# *In-Situ* Structure and Topography of AMPA Receptor Scaffolding Complexes Visualized by CryoET

**DOI:** 10.1101/2024.10.19.619226

**Authors:** Richard G. Held, Jiahao Liang, Luis Esquivies, Yousuf A. Khan, Chuchu Wang, Maia Azubel, Axel T. Brunger

## Abstract

Most synapses in the brain transmit information by the presynaptic release of vesicular glutamate, driving postsynaptic depolarization through AMPA-type glutamate receptors (AMPARs). The nanometer-scale topography of synaptic AMPARs regulates response amplitude by controlling the number of receptors activated by synaptic vesicle fusion. The mechanisms controlling AMPAR topography and their interactions with postsynaptic scaffolding proteins are unclear, as is the spatial relationship between AMPARs and synaptic vesicles. Here, we used cryo-electron tomography to map the molecular topography of AMPARs and visualize their *in-situ* structure. Clustered AMPARs form structured complexes with postsynaptic scaffolding proteins resolved by sub-tomogram averaging. Sub-synaptic topography mapping reveals the presence of AMPAR nanoclusters with exclusion zones beneath synaptic vesicles. Our molecular-resolution maps visualize the predominant information transfer path in the nervous system.

## Main Text

Neuronal synapses detect the release of chemical neurotransmitters using ligand-gated receptors to alter membrane potential and activate downstream signaling events. Glutamatergic synapses, representing ∼80% of synapses in the vertebrate cortex and hippocampus, primarily signal through ionotropic α-amino-3-hydroxy-5-methyl-4-isoxazolepropionic acid receptors (AMPARs) (*1*–*4*). AMPARs are tetrameric ion channels composed of a transmembrane domain, an extracellular ligand-binding domain (LBD), and an extracellular amino-terminal domain (NTD) arranged as a dimer of dimers (*5*–*7*). Glutamatergic signaling through AMPARs is essential for both basal synaptic transmission and dynamic changes in synaptic strength (*i.e*., synaptic plasticity) (*8*). Fusion of a single synaptic vesicle elevates local synaptic cleft glutamate concentration into the low mM range, which rapidly equilibrates to ∼100 μM within ∼80 μs (*9, 10*). AMPARs, however, are activated with EC_50_ values ∼400 μM (*11*–*13*). The result is a steep spatial gradient of receptor activation relative to presynaptic neurotransmitter release sites, with only AMPARs within ∼100 nm of a release site activated (*14*–*17*). Therefore, the molecular topography of AMPARs – their spatial distribution and organization within synapse ultrastructure – is a crucial determinant of synaptic strength and plasticity (*18*–*24*). AMPARs localize to the synapse primarily via interactions between auxiliary Transmembrane AMPA receptor Regulatory Proteins (TARPs) and postsynaptic Membrane-Associated Guanylate Kinases (MAGUKs), most prominently PSD-95 (*25*–*29*). Both AMPARs and PSD-95 form sub-synaptic nanoclusters, tens of nanometers in diameter within individual synapses (*18*–*20, 30*).

These nanoclusters are proposed to act as signaling hotspots, potentiating synaptic response amplitude by increasing receptor density opposite synaptic vesicle fusion sites (*20, 21*). This model is supported by the trans-synaptic alignment of the cytosolic scaffolding proteins of the postsynaptic density (PSD), including PSD-95, and the presynaptic active zone complex (AZ), which controls synaptic vesicle membrane docking and priming to fusion competence (*20, 22, 23, 31*–*34*); Fig. 1A). However, direct visualization of trans-synaptic alignment between docked synaptic vesicles and AMPAR nanoclusters is lacking, as is structural information about the protein complexes that control AMPAR topography. This limits our understanding of how synapses and other cellular compartments organize their signaling pathways.

**Fig. 1.**
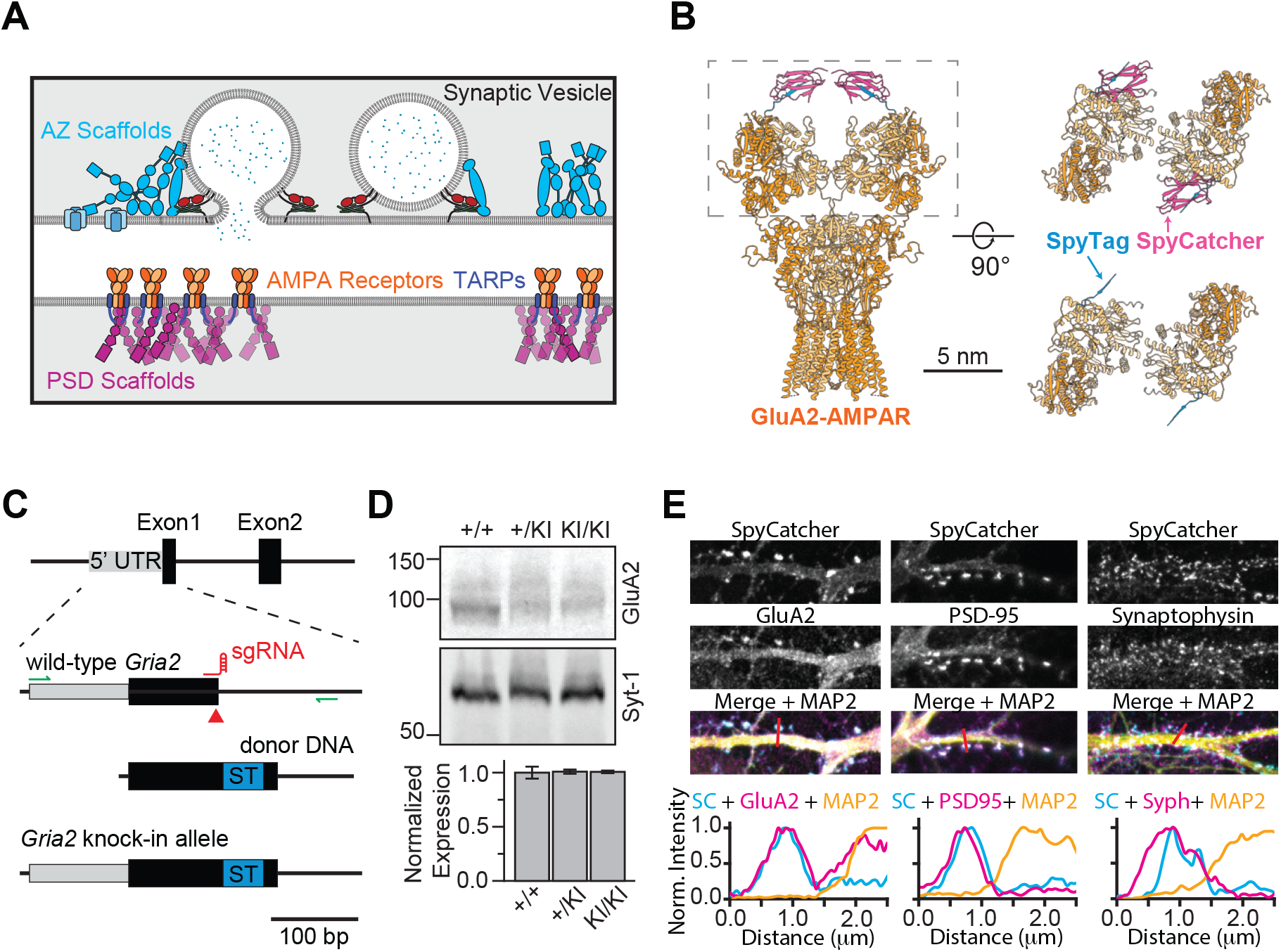
Generation of SpyTag-GluA2 Knock-In Mice. **A)** Schematic of a glutamatergic synapse. Presynaptic vesicles are docked proximal to the plasma membrane by the scaffolds of the active zone (AZ: light blue). Scaffolding proteins of the postsynaptic density (PSD: purple), anchor ionotropic glutamate receptors, including AMPARs (orange) through auxiliary TARP subunits (dark blue). **B)** Model of a SpyTag-GluA2 containing AMPAR (based on PDB 7OCA, 4MLI), labeled with SpyCatcher (GluA2 subunits: light orange; SpyTag: blue; SpyCatcher: magenta). Side-view (left) and top-down views of the boxed region, with and without SpyCatcher (right) are shown. **C)** Knock-in (KI) strategy for generation of SpyTag-GluA2 mice. Double-strand breaks (red arrow) were generated in *Gria2* using sgRNA:Cas9 and repaired with donor DNA containing SpyTag (ST) inserted into the open reading frame of exon 1. The final knock-in allele contains SpyTag at the N-terminal of GluA2, immediately following the signal peptide. **D)** Fluorescent western blots and quantification of GluA2 levels of wild-type (+/+, N=3), heterozygous (+/KI, N=2), and homozygous knock-in (KI/KI, N=3) brain membrane fractions. **E)** Confocal imaging and line-scan analysis of cultured SpyTag-GluA2 knock-in neurons, surface stained with fluorescent SpyCatcher (SC; top panels) followed by fixation and immunostaining against the indicated targets. Line scans were performed over regions indicated by the red line and normalized.

Cryo-electron tomography (cryoET) provides nanometer-scale resolution and the ability to visualize 3D cellular ultrastructure under near-native conditions (*35*). CryoET also enables *in-situ* protein structure determination using sub-tomogram averaging (STA), allowing direct visualization of the relationship between a protein’s structural state and cellular context, such as residency in a sub-synaptic nanocluster (*36*–*39*). Using cryoET, we recently confirmed the presence of trans-synaptically aligned PSD and AZ nanoclusters but, surprisingly, found that membrane-proximal synaptic vesicles are not preferentially aligned to these clusters (*40*). The lack of alignment led us to question whether AMPARs are aligned to membrane-proximal synaptic vesicles, which fuse to release neurotransmitters. While cryoET can, in principle, be used to localize single molecules such as individual AMPARs, in practice, this is limited by the crowded environment of the synapse and the ability to distinguish structurally similar proteins at the resolution of individual tomograms. Therefore, localizing target proteins with high confidence requires specific labels, visible in cryoET reconstructions, or correlative approaches combining fluorescence and electron microscopy (*41*–*45*). Here, we developed a gold-nanoparticle particle (AuNP) labeling strategy for synaptic AMPARs, using a knock-in mouse line to tag AMPARs. Our approach enables the contextual localization and *in-situ* structure determination of these endogenously expressed AMPARs and their binding partners. Our success with this strategy suggests that it can be broadly applied to other target proteins.

### Covalent Labeling of Endogenous GluA2-Containing AMPA Receptors

AMPAR tetramers comprise subunits GluA1-4 in variable stoichiometries and auxiliary subunits, including TARPs and Cornichons (CNIHs) (*7, 27*). While GluA1-4 can form homomeric tetramer assemblies, positions B and D of the tetramer are almost exclusively occupied by the GluA2 subunit in natively purified AMPARs (*7, 46*). To label these GluA2-containing AMPARs (GluA2-AMPARs), we generated a knock-in mouse line using CRISPR-Cas9 and homology-directed repair to introduce the small peptide tag SpyTag into the N-terminus of GluA2, immediately following the endogenous signal peptide (Fig. 1B, C, fig. S1). SpyTag forms an isopeptide bond with its binding partner SpyCatcher, enabling covalent labeling of tagged proteins (*47*). This tag position was previously used to generate knock-in mice containing super-ecliptic pHluorin or AP tag labels without impacting synaptic transmission (*48, 49*). We confirmed successful integration of the SpyTag using nested-PCR and Sanger sequencing (fig. S1C). SpyTag-GluA2 knock-in mice were born in normal Mendelian ratios and could be maintained as homozygous breeding pairs, indicating no loss-of-function due to the introduction of SpyTag (fig. S1A, B).

Homozygous and heterozygous knock-in mice expressed GluA2 at levels indistinguishable from gender-matched wild-type littermates (Fig. 1D). Live-cell surface staining using fluorescently labeled SpyCatcher specifically labeled cultured neurons from knock-in mice with negligible labeling of wild-type controls (fig. S2). To determine if the introduction of SpyTag altered the localization of GluA2-AMPARs, we surface-stained cultured neurons with fluorescent SpyCatcher followed by fixation, permeabilization, and immunostaining with antibodies targeting either the intracellular region of GluA2, PSD-95, or the presynaptic vesicle marker synaptophysin (Fig. 1E). SpyCatcher staining colocalized with GluA2 immunostaining primarily at the ends of dendritic spines, with antibody staining revealing an additional intracellular population of GluA2. PSD-95, which binds AMPARs via interactions with TARP auxiliary subunits, is strongly colocalized with SpyCatcher at spine heads (*26, 50*).

Synaptophysin, which labels presynaptic vesicles in both glutamatergic and GABAergic synapses, also largely colocalized with SpyCatcher staining, though it was slightly offset, in keeping with the localization of vesicles within the presynapse (*51*). These data indicate that adding SpyTag to GluA2 allows the specific, extracellular, and covalent labeling of endogenously expressed GluA2-AMPARs without compromising receptor function (*49*), expression, or localization.

### Cryo-Electron Tomography Localizes Gold Nanoparticle Labeled AMPARs

Next, we imaged endogenous GluA2-AMPARs with high precision in 3D within synaptic ultrastructure. To achieve this, we used SpyCatcher to functionalize AuNPs (Fig. 2A). AuNPs exhibit high contrast in cryoEM (Fig. 2B) and have been previously used to label cell surface proteins (*41, 52, 53*). We synthesized thiol-protected AuNPs with uniform diameter of ∼2.5 nm measured by TEM and dynamic light scattering (DLS) (fig. S4A, B). AuNPs were functionalized 1:1 with SpyCatcher via place-exchange reactions, followed by passivation of the remaining active sites on the protected layer of the AuNP (fig. S3A-D). The particle diameter of SpyCatcher-AuNP is ∼4 nm as measured by DLS (fig. S4C). Conjugation of AuNP to SpyCatcher did not disrupt its ability to bind with SpyTag-MBP in native gel shift assays (Fig. 2C, fig. S4D). This binding was specific, as MBP alone did not interact with SpyCatcher-AuNPs. Complete quenching of AuNP reactivity and the stability of passivation with glutathione was further confirmed (fig. S3C, D). Altogether, these experiments confirm the proper function and specificity of our SpyCatcher-functionalized AuNPs.

**Fig. 2.**
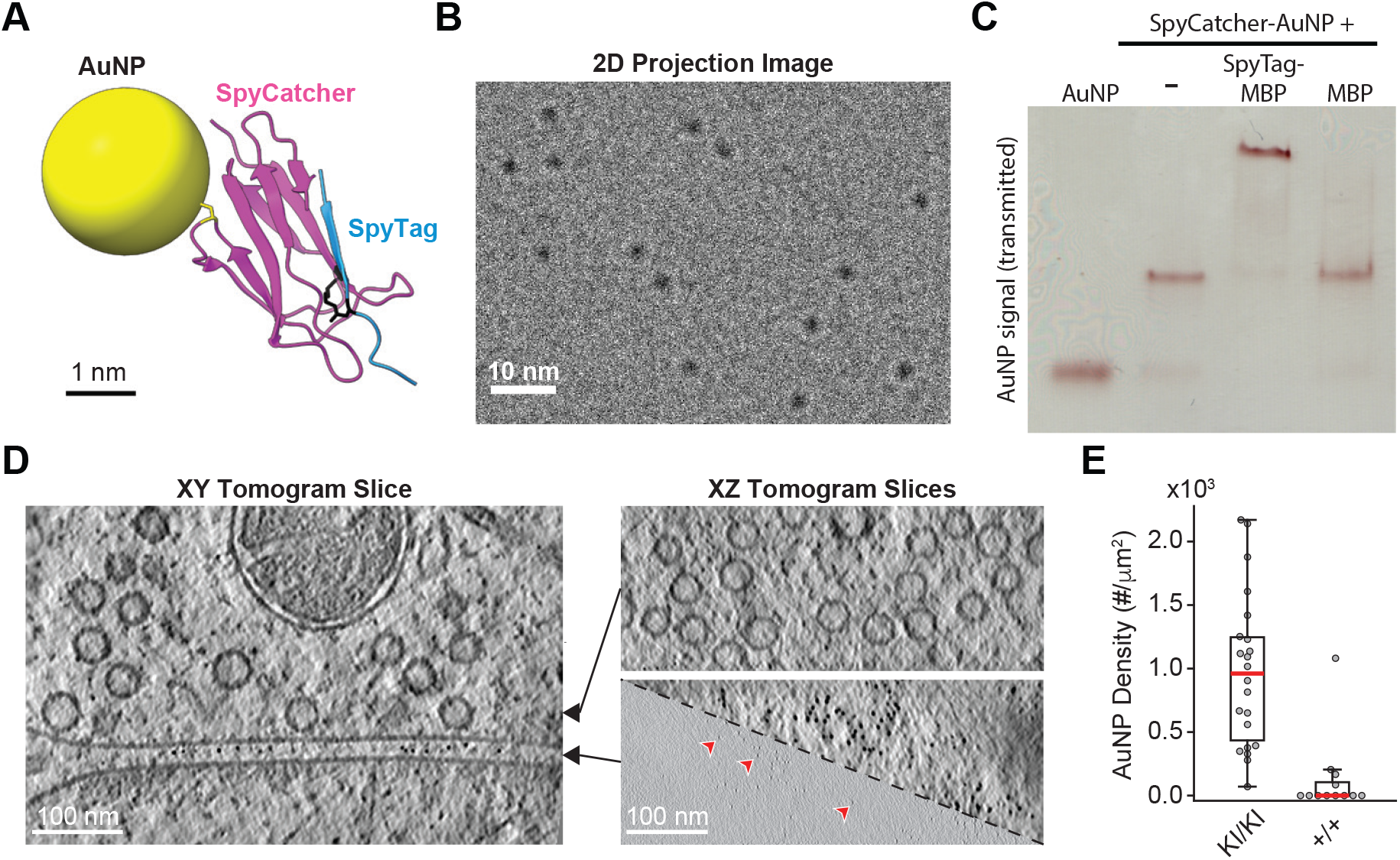
SpyCatcher-AuNP Labeling of SpyTag-GluA2 AMPARs. **A)** Scale model of SpyCatcher-functionalized AuNPs (based on PDB:4MLI). SpyCatcher is covalently linked to the AuNP via the sulfhydryl group of cysteine 49 (yellow stick models). SpyTag forms an isopeptide bond between an Asn residue and SpyCatcher Lys 31 (black stick models). **B)** Cryo-TEM projection images of AuNPs at a defocus of 1.5 μm. **C)** Native gel shift assay showing unconjugated AuNPs, 1:1 SpyCatcher-AuNPs, and SpyCatcher-AuNPs in the presence of excess SpyTag-MBP or MBP. **D)** Slices through a tomogram of a SpyTag-GluA2 knock-in synapse, labeled with SpyCatcher-AuNPs, reconstructed at a pixel size of 13.88Å/px (bin-8) and processed with IsoNet (*66*). Right panels show XZ slices at the plane indicated by the arrows. The top right slice through the membrane-proximal vesicle layer is at bin-8 and processed with IsoNet. The bottom right slice shows the synaptic cleft layer at both bin-8 and bin-2 (3.47Å/px) without IsoNet processing. Red arrowheads point to AuNPs, which show high contrast signal even in unprocessed bin-2 tomograms. **E)** Quantification of AuNP density as the number of AuNP localizations per μm^2^ of PSD membrane area in SpyTag-GluA2 knock-in (KI/KI, N=22) and wild-type (+/+, N=12) synapses. Mann Whitney U tests were used to test for significance (***; p < 0.001).

Next, we grew primary hippocampal neurons from SpyTag-GluA2 knock-in mice directly on EM grids and stained with SpyCatcher-AuNPs. After washing, SpyCatcher-AuNP labeled neurons were plunge-frozen in liquid ethane, and cryo-focused ion beam (FIB) milling was used to prepare 150-250 nm thick lamellae of fasciculated dendritic and axonal processes containing synapses (*40*). We used these lamellae samples for cryoET, targeting synapses for tilt series data collection and tomogram reconstruction. AuNPs were clearly visible within the synaptic cleft and were readily distinguishable from the surrounding biological material by their high contrast even at low voxel bin values (Fig. 2D). SpyTag-GluA2 knock-in synapses contained abundant AuNPs with a median density of 961 AuNPs/μm^2^ within the PSD membrane (Fig. 2E). As a control, the addition of SpyCatcher-AuNPs to wild-type synapses showed a near-complete loss of AuNP labeling, indicating specific labeling of SpyTag-GluA2 in knock-in neurons.

To assess the AMPAR labeling efficiency, we performed competitive binding experiments using fluorescence microscopy, first labeling with SpyCatcher-AuNP, then wash-out and subsequent labeling with fluorescent SpyCatcher (fig. S2A, B). Under these conditions, ∼65% of the fluorescence signal is lost compared to fluorescent labeling of knock-in neurons without pre-incubation with SpyCatcher-AuNP. The remaining signal reflects a mix of AMPARs newly trafficked to the plasma membrane after the removal of SpyCatcher-AuNP and those unlabeled during the initial incubation. Notably, some synapses appeared to have a complete loss of fluorescent labeling, indicating that newly trafficked receptors likely account for most of the remaining signal (fig. S2A, B). Overall, our SpyTag-GluA2/SpyCatcher-AuNP system enables sensitive and specific labeling of endogenously expressed GluA2-AMPARs in the crowded environment of the synaptic cleft.

### Nanometer-Scale 3D Spatial Statistics of GluA2-AMPARs

Using the 3D localizations of AuNP-labeled GluA2-AMPARs, we analyzed the spatial statistics of AuNP point patterns and, to assess significance, compared our observed localization data to simulated random data (Fig. 3A). We first analyzed the nearest-neighbor distances (NND) between AuNP localizations. There were multiple prominent peaks in the NND histogram, which were fit with a three-Gaussian function model with peaks at 6.02, 12.18, and 23.42 nm (Fig. 3B). We modeled the expected AuNP positions and NND with GluA2 occupying the B and D positions of the AMPAR tetramer using published high-resolution structures of AMPARs (Fig. 3C, fig. S5). Modeled NNDs were bimodal, with average values of 5.68 +/-1.32 nm (15 of 20 Au-NP modeled structures) and 12.26 +/-3.13 nm (5 of 20 AuNP-modeled structures), consistent with the observed peaks in our NND histogram (Table S1). Resolving two NND peaks separated by ∼6 nm, consistent with published AMPAR structures, suggests that this approach can provide information about the conformational state of individual proteins in a cellular context. Moreover, 88% of AuNPs had a nearest-neighbor within 17 nm – the maximum inter-AuNP distance in our models – supporting the notion that our labeling is close to saturation for those AMPARs present on the membrane during the labeling period, in approximate agreement with our competition experiments (fig. S2A, B). Since both GluA2 subunits of most AMPARs were labeled with SpyCatcher-AuNPs and labeling was specific in-vitro and in cells, we refer to AuNP localizations as GluA2-AMPARs going forward.

**Fig. 3.**
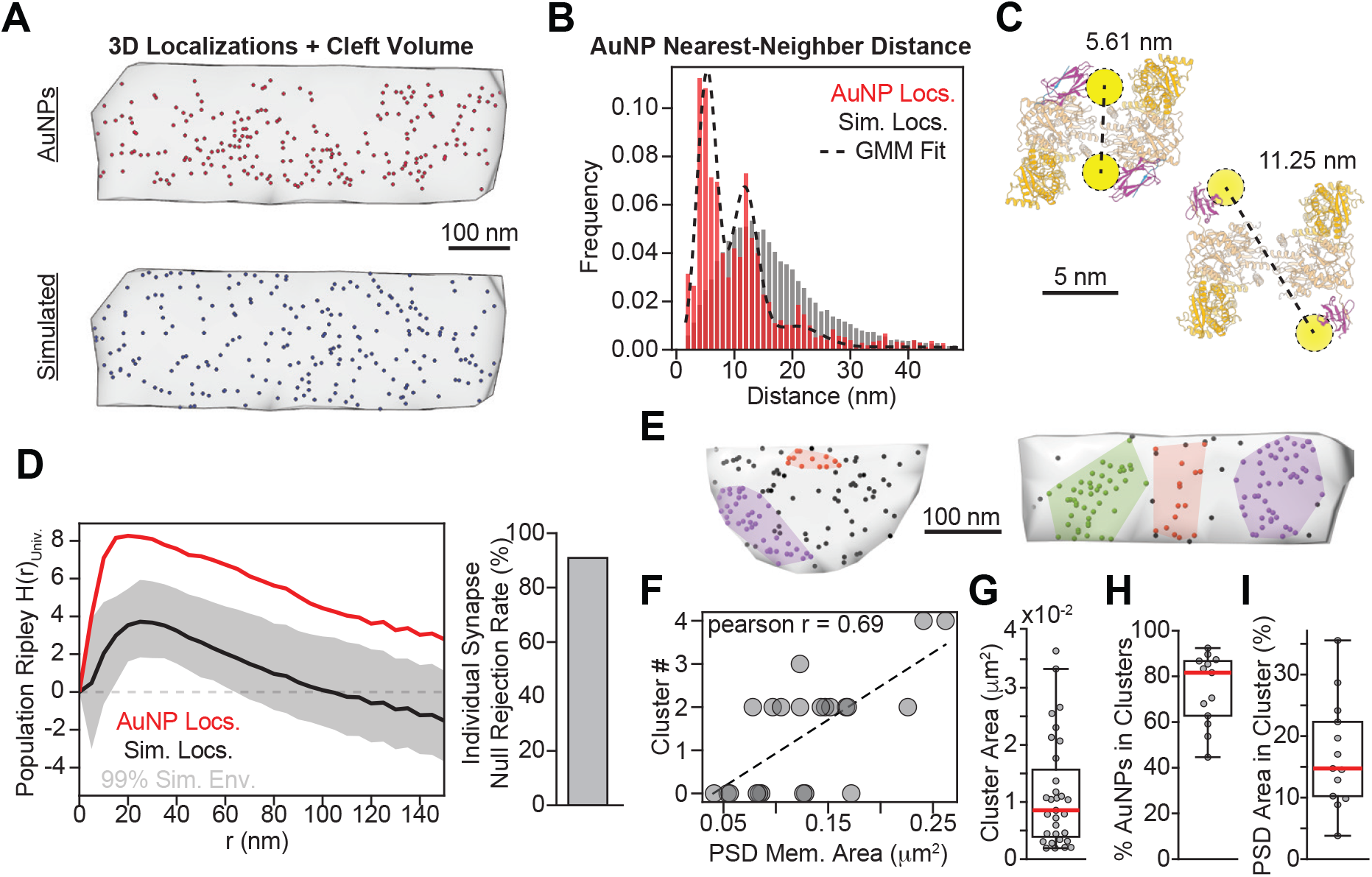
Spatial Statistics of Endogenous GluA2-AMPARs. **A)** 3D localizations of AuNPs (top) and simulated random data (bottom) within the synaptic cleft (grey volume). **B)** Nearest-neighbor distance (NND) values between AuNPs (red) compared to simulated data (grey). The data histogram was fit with a Gaussian mixture model (gmm; dashed line). **C)** Models of SpyTag-GluA2 AMPARs, labeled with SpyCatcher-AuNPs based on PDB: 7LDD (top left), 8SS8 (bottom right), and 4MLI (SpyCatcher/Tag). **D)** Quantification of AuNP clustering by Ripley H analysis. The black line and grey band represent the mean and 99% simulation envelope of simulated random data (N=1100). The red line represents the pooled mean of AuNP localization data (N=22). Significance was assessed by MAD tests (***; p<0.001). The right panel shows the results of MAD tests conducted on individual synapses and their respective simulation data (N=50/synapse tomogram). **E)** Examples of AuNP clusters identified by HDBSCAN, color-coded, and outlined by their convex hull. **F)** Relationship between PSD membrane area and the number of clusters identified by HDBSCAN. **G)** Area of clusters identified by HDBSCAN (N=31 clusters). **H)** Percentage of AuNP localizations found within clusters for synapses with identified clusters (N=13 synapses). **I)** Percentage of the PSD membrane area covered by clusters for synapses with identified clusters (N=13 synapses).

We next analyzed GluA2-AMPAR clustering using Ripley H functions, which quantify the number of neighboring AuNPs within a given radius from each particle (Fig. 3D). Population analysis of all synapses in the dataset revealed that GluA2-AMPARs are significantly clustered across spatial scales, consistent with visual inspection of point patterns compared to simulated random data. When each synapse was analyzed individually, the null hypothesis (*i.e*., random placement of the same number of AuNPs in the synaptic cleft) was rejected in 91% (20/22 synapses) of synapses analyzed (Fig. 3D). These data support the notion that GluA2-AMPARs are clustered at the synapse, consistent with super-resolution fluorescence microscopy experiments (*18, 19, 54, 23, 49, 55*). To visualize discrete GluA2-AMPAR clusters, we performed hierarchical density-based spatial clustering analysis (HDBSCAN) (*56*) to assign cluster labels to AuNP localizations (Fig. 3E). Cluster numbers per synapse correlated with the area of the PSD membrane contained within the lamella and ranged from zero to four clusters per synapse (Fig. 3F). The median cluster area was 0.0085 μm^2^ (Fig. 3G), consistent with previous estimates of AMPAR nanocluster size (*18, 19*). For those synapses containing clusters, a median of 81.6% of AuNP labels were localized to a cluster, and 15.7% of the total PSD membrane area was occupied by AMPAR clusters (Fig. 3H, I).

### Contextual Localization of GluA2-AMPARs

The spatial organization of synaptic AMPARs and their topography relative to neurotransmitter release sites are key determinants of the amplitude of the postsynaptic response to neurotransmitter release (*17*–*20, 22, 23*). We sought to map the molecular topography of synaptic AMPARs by relating specific AuNP localization data to ultrastructural information provided by cryoET, going beyond our previous study without GluA2-AMPAR labels (*40*). For each synapse tomogram, synaptic vesicles less than 10 nm from the plasma membrane (*i.e*., membrane-proximal) and PSD protein density were segmented and visualized in relation to AuNP localizations (Fig. 4A). The clustered point patterns of GluA2-AMPARs often appeared to have visible holes which, intriguingly, are occupied by membrane-proximal synaptic vesicles (fig. S6). We refer to this organization as an excluded topography (*57, 58*).

**Fig. 4.**
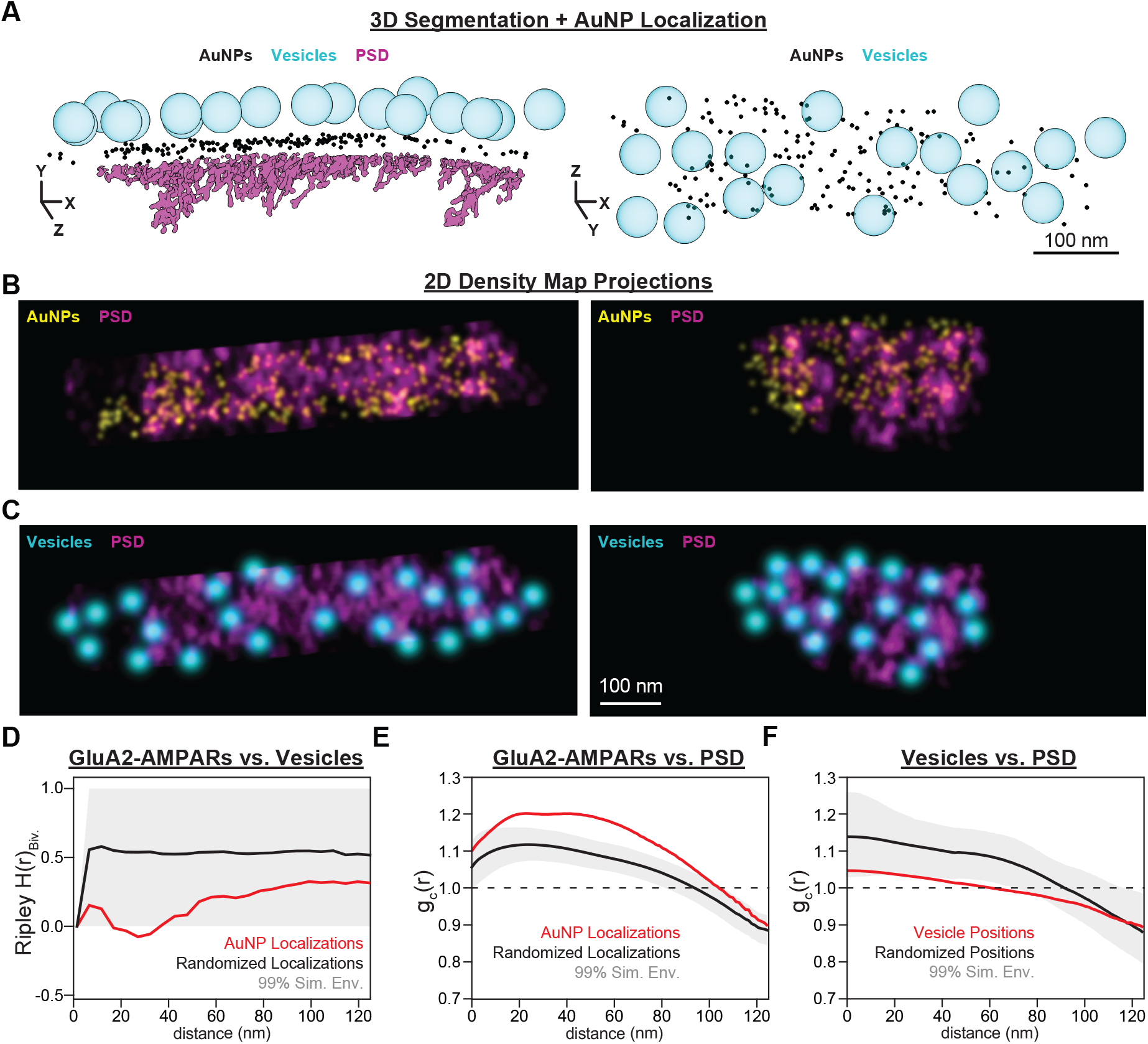
Molecular Topography of Synaptic GluA2-AMPARs. **A)** Volume rendering of a segmented synapse tomogram showing membrane-proximal synaptic vesicles (cyan), AuNPs (black), and PSD protein density (magenta). AuNPs are shown at a diameter of 4 nm. (Right) top-down view showing membrane-proximal synaptic vesicles (cyan) and AuNP localizations (black). **B)** Overlaid projections of PSD (magenta) and AuNP (yellow) density maps. **C)** Overlaid projections of PSD (magenta) and synaptic vesicle (cyan) density maps. **D)** Standardized bivariate Ripley H analysis of the relationship between synaptic vesicle and AuNP localizations. The red line shows the mean and standard error of AuNP localization data from all synapses (N=21). The black line and gray boundary show the mean of simulated data and 99% simulation envelope (N=2310). Significance was assessed using MAD testing (*, p<0.05). **E)** The normalized paired cross-correlation, g_c_(r), between AuNP localizations and the local density of PSD material. The red line shows the mean of all synapses (N=22 synapses). The black line and gray boundary show the mean of simulated AuNP localization data and 99% simulation envelope (N=5000), respectively. Significance was assessed using MAD testing (***, p<0.001). **F)** g_c_(r), between synaptic vesicle positions and the local density of PSD material (N=21 synapses, 5000 simulations). Significance was assessed using MAD testing (*, p<0.05).

Bivariate Ripley H function analysis of the entire population of synapses (Fig. 4D) revealed a significant decrease in the number of AuNP localizations surrounding synaptic vesicles, which extend outside the 99% simulation envelope in the range of 20-40 nm, approximately corresponding to the average radius of a synaptic vesicle. Analysis of individual synapses revealed that, while all synapses in the dataset exhibited decreased GluA2-AMPAR numbers surrounding synaptic vesicles, only 62% (13/21 synapses) reached the significance threshold and could be said to adopt an excluded topography (fig. S7A, B). Synapses without significant GluA2-AMPAR exclusion had significantly fewer nanoclusters identified by HDBSCAN, indicating that increased spatial clustering of GluA2-AMPARs corresponds with excluded topography relative to membrane-proximal synaptic vesicles (fig. S7C-F). Indeed, at a qualitative level, synaptic vesicles appeared to decorate the perimeter of GluA2-AMPAR nanoclusters (fig. S6). Together, GluA2-AMPARs in the synaptic cleft are preferentially excluded from the region opposite membrane-proximal synaptic vesicles.

We next asked whether the density of GluA2-AMPARs was related to the local density of the PSD. We converted PSD segmentation and AuNP localizations into voxel-based local density maps and analyzed the relationship between the two as a normalized paired cross-correlation function, g_c_(r) (Fig. 4B). GluA2-AMPAR density was significantly correlated with the PSD density – which was non-uniform as previously described (*18, 40*) – with correlation values peaking at shift ranges between 20-50 nm (Fig. 4E). These data are consistent with the AMPAR scaffolding function of the PSD and suggest that most SpyCatcher-AuNP labeled GluA2-AMPAR complexes are anchored at the synapse through interactions with scaffolding proteins such as PSD-95. Moreover, overlaid local density maps and paired cross-correlation analysis of the PSD and synaptic vesicle positions showed synaptic vesicles occupy low-density regions of the PSD and surrounding high-density peaks (Fig. 4C, 4F).

### Structure of AMPAR Scaffolding Complexes

Given the correlation between GluA2-AMPARs and PSD density, we asked whether stereotyped interactions between AMPARs and the scaffolding proteins of the PSD might occur. Using sub-tomogram averaging (STA), we first determined the structure of labeled GluA2-AMPARs at a resolution of 31 Å by focusing on the extra-cellular domains (ECD) (Fig. 5A-B, fig. S9); suppression of AuNP and membrane signal was essential for the initial alignments (fig. S8) (Methods). This AMPAR-ECD map suggests an AMPAR dimer-of-dimer quaternary structure, including the two-layer LBD and NTD structure with apparent domain swapping between layers. Derived from 618 sub-tomograms, the map quality highlights the advantages of high-confidence particle picking using specific labels. The map could accommodate published structures of natively purified AMPARs (*59*), with additional density, likely contributed by SpyCatcher-AuNPs, extending from the B/D subunits.

**Fig. 5.**
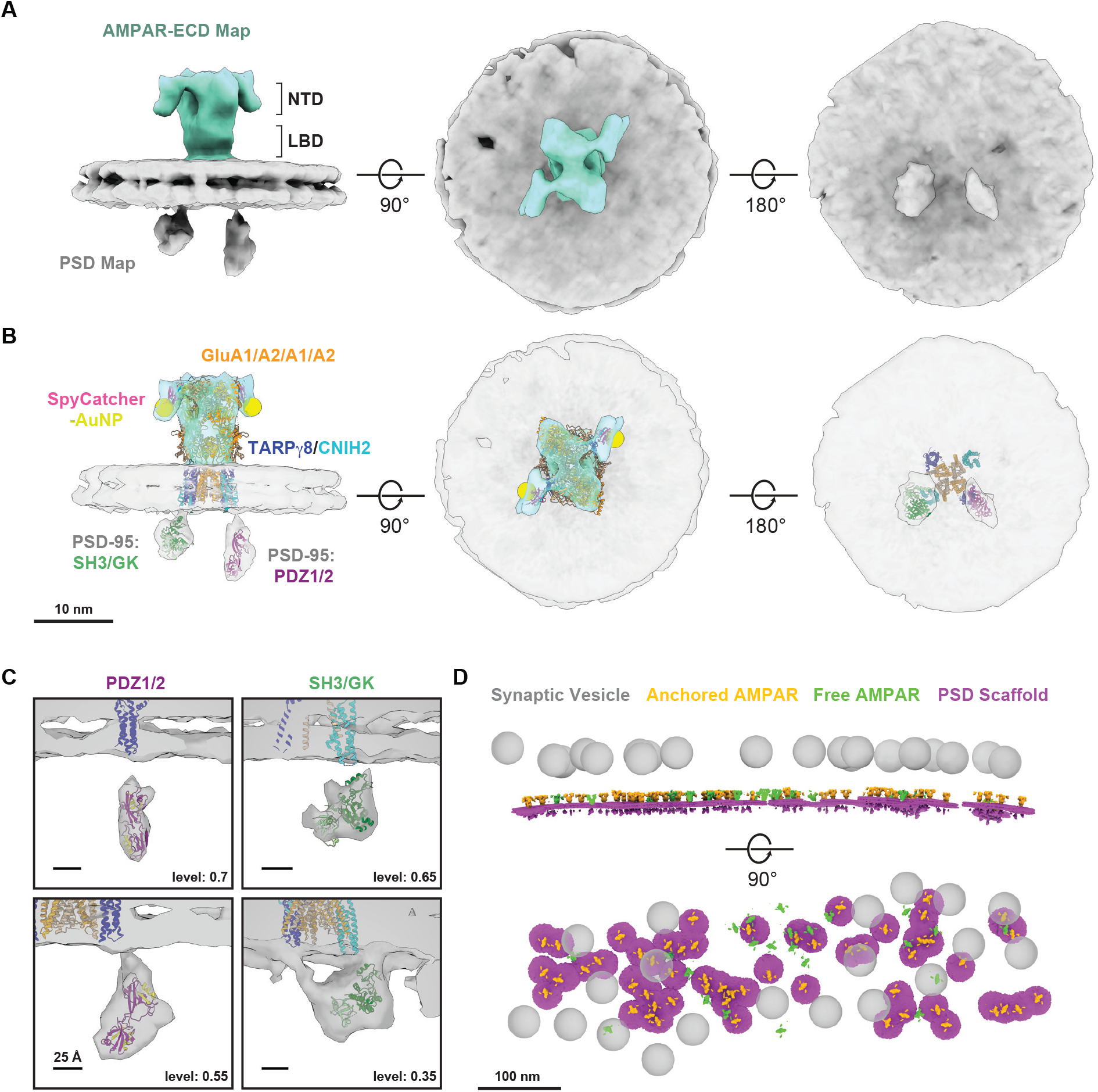
STA of AMPAR/PSD Scaffolding Complexes. **A)** Composite sub-tomogram average maps of the GluA2-AMPAR extracellular domains (AMPAR-ECD Map, teal) and AMPAR-associated scaffolding proteins of the PSD (PSD Map, grey). **B)** Model of the AMPAR/PSD scaffolding complex assembled by rigid body docking high-resolution structures of native AMPARs (PDB-7LDD: GluA1/A2, TARP-γ8, CNIH2; PDB-4MLI: SpyCatcher/Tag; PDB-3ZRT: PSD-95 PDZ1/2 domain, PDB-1JXO: PSD-95 SH3/GK. **C)** Close up views of two main PSD scaffold densities at different contour levels and orientations. Left: The elongated density with the PDZ1/2 domains of PSD-95 docked and apparent connecting density to TARP-γ8. The binding groove of each PDZ domain is highlighted in yellow. Right: The globular density with the SH3/GK domain of PSD-95 docked, apparently bridging TARP-γ8 and CNIH2. **D)** Representative AMPAR-ECD (yellow) and PSD (purple) STA maps back-plotted to their tomogram positions and displayed along with membrane-proximal synaptic vesicles (cyan).

We then shifted the focus of the analysis to the PSD region directly beneath each receptor. Starting from the alignments from the AMPAR-ECD map, 3D classification and refinement yielded a map of the PSD at 28 Å from 460 sub-tomograms (Fig. 5A-B, fig. S10). Two prominent and ordered postsynaptic protein densities were apparent, one extended and one globular, both in apparent contact with the auxiliary subunits of the AMPAR complex, which could be docked in the transmembrane region of the map (Fig. 5A-C, fig. S11A). The extended density accommodated crystal structures of the tandem PDZ1/2 domains of the PSD-95 (*60*).

PSD-95 binds to TARPs, is the most abundant MAGUK, and is among the most abundant proteins in the PSD (*26, 61, 27*). These extended densities may be related to extended features observed in lower-resolution tomograms from high-pressure frozen, freeze substituted samples that are dramatically reduced by genetic knock-down of MAGUKs (*28*). Our STA map places PDZ1/2 directly beneath the TARP-γ8 in the docked model (Fig. 5C). The globular density appeared to primarily contact the membrane near the CNIH2 auxiliary subunit, but also showed an extended tail connecting to the second TARP-γ8 at lower surface contour levels (Fig. 5C, bottom right panel). This density accommodates the SH3-GK domains of PSD-95 (Fig. 5C) (*62*). Alternatively, this density could also fit a second PDZ1/2 pair, albeit the shape of the density is more consistent with a SH3-GK domain (fig. S11B). In support of our proposed fit, the SH3-GK also binds to TARPs and stabilizes PSD-95 at the synapse (*63, 64*). At lower contour values, additional densities were apparent and fit well with the folded domains of PSD-95, which are separated by long flexible linkers (fig. S11C). Together, our STA maps revealed an extensive network of interactions between AMPARs and PSD molecules that is sufficiently well-ordered to emerge in these averaged maps.

We plotted the AMPAR and PSD maps back in the coordinate system of our tomograms along with membrane-proximal synaptic vesicles (Fig. 5D). Consistent with density correlation and point-pattern analysis (Fig. 4), these topography maps revealed clusters of AMPARs associated with PSD scaffolds and excluded from areas under synaptic vesicles, which decorated their perimeter. AMPARs without associated PSD scaffolds were largely isolated from or at the edges of these clusters. GluA2-AMPARs residing in nanoclusters (∼82% of AuNP labels, Fig. 3H) are, therefore, preferentially engaged in surprisingly structured interactions with the PSD (74% of AMPAR sub-tomograms, fig. S10).

In summary, we functionalized AuNPs with SpyCatcher, specifically and covalently labeled and localized endogenously expressed GluA2-AMPARs using SpyTag-GluA2 knock-in mice, and visualized their *in-situ* structure and topography with cryoET. Our modular approach uses labels that are smaller than conventionally used antibody-based methods. Our small labels minimize localization errors and efficiently access crowded extracellular spaces like the synaptic cleft. Using this labeling system, we resolved individual subunits of single AMPARs and contextualized single-molecule localization with high-resolution cellular ultrastructure. We established high-confidence particle picks that enable *in-situ* structure determination of both target and associated non-target proteins. Our STA maps reveal surprisingly structured complexes of AMPARs with PSD scaffolding proteins, most likely involving the MAGUK PSD-95 and TARP auxiliary subunits. The PSD, therefore, contains a stereotyped local structure amenable to *in-situ* structure determination. Thus, around AMPARs, PSD molecules do not engage in highly transient interactions, as in a protein condensate (*64, 65*), but participate in a well-defined network that likely contributes to AMPAR stabilization and clustering. Sub-synaptic nanoclusters of GluA2-AMPARs are not directly aligned with membrane-proximal synaptic vesicles. Instead, the volume of the synaptic cleft directly beneath a membrane-proximal synaptic vesicle appears to behave as an AMPAR exclusion zone, resulting in an offset between clustered GluA2-AMPARs and synaptic vesicles. This observed excluded topography is reminiscent of observations for synaptic vesicles and presynaptic voltage-gated calcium channels (*57, 58*). We speculate that the molecular topography of the synapse is driven in part by spatial constraints imposed by the bulky pre- and postsynaptic scaffolding complexes of the AZ and PSD, which act via a “push-and-pull” mechanism to physically occlude relatively large synaptic vesicles from high-density regions while clustering key synaptic proteins like AMPARs. In conclusion, we envision creating molecularly precise maps of protein topography and atomic models of complex networks of protein interactions – even in relatively unstructured cellular environments – by building out models of higher-order complexes stepwise from structures of labeled targets.

## Acknowledgments

This article is subject to HHMI’s Open Access to Publications policy. HHMI laboratory heads have previously granted a non-exclusive CC BY 4.0 license to the public and a sublicensable license to HHMI in their research articles. Pursuant to those licenses, the author-accepted manuscript of this article can be made freely available under a CC BY 4.0 license immediately upon publication. The project described was supported, in part, by Award Number 1S10OD010580 from the National Center for Research Resources (NCRR). Its contents are solely the responsibility of the authors and do not necessarily represent the official views of the NCRR or the National Institutes of Health. We also acknowledge funding from the Phil and Penny Knight Initiative for Brain Resilience at the Wu Tsai Neurosciences Institute, Stanford University. Some of this work was performed at the Stanford-SLAC CryoET Specimen Preparation Center (SCSC) which is supported by the National Institutes of Health Common Fund’s Transformative High Resolution Cryoelectron Microscopy program (SCSC: U24GM139166). The authors also acknowledge the Stanford University Cryo-electron Microscopy Center (cEMc) for equipment access, the Stanford Transgenic, Knockout, and Tumor Model Center for assistance with generating mice, and the lab of Liang Feng for assistance with flow cytometry experiments. The authors also thank Allister Burt and Dimitry Tegunov for their help in implementing Warp2.0 for Linux on our computing cluster, and Reza Paraan, Saugat Kanel, and Shawn Zheng for helpful discussions on image processing of membranes.

## Funding

Funding was provided by the US National Institutes of Health (NIH; RO1MH63105 to A.T.B.)

## Author contributions

Conceptualization: RGH, MA, ATB

Methodology: RGH, MA, ATB

Investigation: RGH, JL, LE, YAK, CW

Visualization: RGH, JL

Funding acquisition: ATB

Project administration: RGH, ATB

Supervision: ATB

Writing – original draft: RGH, JL, ATB

Writing – review & editing: RGH, JL, LE, YAK, CW, MA, ATB

## Competing interests

Authors declare that they have no competing interests.

## Data and materials availability

All analysis code will be available on the Brunger Lab GitHub page (https://github.com/brungerlab). Sub-tomogram average maps, reconstructed tomograms, and raw tilt series data will be deposited to EMDB and EMPIAR.

## Supplementary Materials

Materials and Methods

Figs. S1 to S11

Tables S1

References (*1-16*)

Movies S1 to S2

## Notes

### Competing Interest Statement

The authors have declared no competing interest.

### Summary of Updates

Added ORCID for one of the authors, fixed minor typos in the manuscript, added movies S1, S2.

