## Supplementary Material for "*In-Situ* Structure and Topography of AMPA Receptor Scaffolding Complexes Visualized by CryoET"

**The PDF file includes:**

Materials and Methods
Figs. S1 to S11
Tables S1
References

**Other Supplementary Materials for this manuscript include the following:**

Movies S1 to S2

### Materials and Methods

Generation of SpyTag-GluA2 Knock-In Mouse

All animal experiments were performed in accordance with Stanford APLAC institutional guidelines under protocol #29981. SpyTag-GluA2 knock-in mice were generated using CRISPR-Cas9 and homology-directed repair targeting exon 1 of the Gria2 gene (chr3:80,709,554-80,710,174; mouse genome assembly GRCm39/mm39). 10 ng/µl sgRNA (CTAACAGCATACAGATAGGT) was mixed with 30 ng/µl purified Cas9 protein at room temperature for ten minutes, followed by addition of single-stranded donor DNA (CTACATGACATGAATTAATTTGCACACTGTGAACAAAGATTACCACCAAATTATGC ACAATTCAAAGGCATAACACAAAAGGTACCTACCTATTTGTATGCTCTTGTATCTCT TGTAGGCGTCCACCATCACGATGTGGGGCACGCCTCTCGAGCCACCGCCACCACTAG TGTTAGAAGAGACACCAAAAATCAGTCCCCATAAAACAGGAGAAAGGAGGACAGA AATATGCATAATCTTTTGCATTTCCAAGAAAAGTAGAGCATCCAC) to 10 ng/µl. 176 embryos from C57BL6 mice were injected with this mixture and implanted into surrogate mothers, resulting in 23 pups and six positive founder mice. PCR was used to confirm the genotype of each founder using oligos outside the dDNA sequence (TGCTTCAGCCTAAGAAAATCATG and GGAGGAAAGGGAAACGAGGG; wild-type band = 302 bp, knock-in band: 371 bp). PCR products were purified by gel extraction and Sanger sequencing was performed using nested primers to confirm correct donor integration. Positive founders were maintained as separate lines during three generations of back-crossing to wild-type C57BL6 mice. After back-crossing, two of the original founder lines, which carried identical mutations, were crossed to maintain a single knock-in line, which was bred to homozygosity and used for subsequent experiments.

Neuronal Culture Preparation

Primary cultures of mouse hippocampal neurons were prepared as previously described (*1*). For cryo-electron tomography experiments, gold 200 mesh Quantifoil R2/2 or R1/4 grids were glow discharged (15 mA, 45 seconds) and placed under UV illumination for 30 minutes. Grids were placed in 35mm glass-bottom dished under a bubble of Matrigel for 1 hour at 37 C before use. For fluorescence microscopy experiments, autoclaved glass coverslips (#1.5) were placed in 24-well plates and coated with a bubble of Matrigel for 1 hour at 37 C before use. Postnatal day 0 (P0) pups of either SpyTag-GluA2 knock-in or wild-type C57BL6 mice were anesthetized on ice and the hippocampus dissected out. Hippocampi were incubated in a papain solution (10 mL Hank's Balanced Salt Solution (HBSS), 10 uL 0.5 M EDTA pH 8.0, 10 uL 1 M CaCl2, 100 uL papain, 100 uL DNAse I) for 15 minutes at 37 C, followed by washes in HBSS and re-suspension in 1 mL per pup of plating medium (Mimimum Essential Medium (MEM) with 0.5% glucose, 0.02% NaHCO3, 0.1 mg/mL transferrin, 10% Fetal Select bovine serum, 2 mM L-glutamine, and 25 mg/mL insulin). After resuspension, cells were dissociated by gentle trituration to generate a single-cell suspension. This cell suspension was bubbled on top of either grids or glass coverslips for 45 minutes, followed by flooding of the entire dish with plating medium. After one day in-vitro (DIV1), the medium was exchanged to growth medium composed of MEM with 0.5% glucose, 0.02% NaHCO3, 0.1 mg/mL transferrin, 5% Fetal Bovine Serum, 2% B-27 supplement, and 0.5 mM L-glutamine. At DIV3-5, half the medium was exchanged to a growth medium supplemented with 4 μM Cytosine b-D-arabinofuranoside (AraC). Cultures were maintained until DIV14-18 before vitrification.

### Western blotting

To quantify GluA2 expression levels by western blotting, three sets of gender-matched littermates from het x het SpyTag-GluA2 knock-in mating pairs were used to prepare brain membrane fractions. In one set, the heterozygous animal had to be euthanized due to injury before sample processing and was not included. Animals were anesthetized using isoflurane, followed by decapitation and removal of brain tissue. Each brain was independently homogenized using a dounce homogenizer in a homogenizing buffer containing 320 mM sucrose, 4 mM HEPES pH 7.4, 1 mM PMSF, and 1mg/mL Pepstatin. Large cellular debris was pelleted by centrifugation at 3500 RPM for 10 minutes using a TA20 rotor. The supernatant from this spin was collected and centrifuged at 100,000 g on a 70Ti rotor for 45 minutes to pellet the membrane fraction. Each pellet was resuspended in a buffer containing 20 mM Tris, 150 mM NaCl, and 10 mM $CaCl_2$ at pH 8. Protein concentration was measured using BCA assays and normalized to 10 mg/mL for each sample. Samples were flash-frozen in liquid nitrogen and stored at -80 C until use.

Western blotting was performed using standard protocols. 50 ug protein from each brain was solubilized in 1X LDS sample buffer for 20 minutes at room temperature (without boiling). Samples were run on Any KD Tris-glycine gels in SDS/Tris/glycine running buffer, then transferred to nitrocellulose membranes using an iBlot3 transfer system. Nitrocellulose membranes were blocked with BioRad EveryBlot blocking buffer for 30 minutes at room temperature, then incubated in primary antibody, diluted 1:1000 in blocking buffer at 4C overnight. Primary antibodies used included: rabbit polyclonal anti-GluA2 (Synaptic Systems Cat# 182 103, RRID:AB_2113732) and mouse monoclonal anti-synaptotagmin-1 (Synaptic Systems Cat# 105 011, RRID:AB_887832). Samples were washed 3-4 times in blocking buffer for 10 minutes then incubated in fluorescently labeled secondary antibody for two hours at room temperature, protected from light. Secondary antibodies used included: goat polyclonal anti-mouse IgG2a Alexa Fluor 488 (Thermo Fisher Scientific Cat# A-21131, RRID:AB_2535771) and goat polyclonal anti-rabbit Alexa 594 (Thermo Fisher Scientific Cat# A-11037, RRID:AB_2534095). Membranes were imaged using an Thermo Fisher iBright imaging system and analyzed in FIJI/ImageJ. GluA2 expression was quantified by first normalizing the integrated density of the GluA2 band to the corresponding synaptotagmin-1 band, then normalizing each lane to the average of all three wild-type lanes.

### Protein Purification

SpyCatcher: pDEST14 SpyCatcher003 S49C was a gift from Mark Howarth (Addgene plasmid # 133448 ; http://n2t.net/addgene:133448 ; RRID:Addgene_133448). The plasmid was transformed into BL21 (DE3) *E. coli* and single colonies were picked and grown overnight to saturation in LB medium and 100 ug/mL ampicillin at 37 C. The following day, starter cultures were used to inoculate 1L flasks of LB + 0.8% (w/v) glucose and 100 ug/mL ampicillin at a 1/100 dilution and grown at 37 C with shaking at 200 RPM until reaching an $OD_{600}$ of 0.6. Cultures were induced with 420 μM Isopropyl β-D-1-thiogalactopyranoside (IPTG) and moved to 30C for 6 hours with shaking at 250 RPM. Cells were pelleted by centrifugation at 5000 g and lysed using a tip sonicator (60% power for 2x 5 minutes, 3 seconds on, 9 seconds off) in buffer containing 50 mM Tris-HCl, 300 mM NaCl, pH 7.8 and protease inhibitors (1 mM PMSF and 1mg/mL Pepstatin). Lysates were centrifuged at 100,000 g for 30 minutes at 4 C, and the supernatant was incubated with 1.5 mL Ni-NTA resin per liter of culture. The bound resin was washed with ten bead volumes of buffer containing 50 mM Tris-HCl, 300 mM NaCl, pH 7.8, and 20 mM imidazole, followed by elution with buffer containing 50 mM Tris-HCl, 300 mM NaCl, pH 7.8, and 250 mM imidazole. The sample was concentrated and then dialyzed overnight at 4C to PBS in a 3.5 MWCO cassette in the presence of 1.5 mg/mL TEV protease to remove the His-tag. Following TEV cleavage, samples were treated with 5 mM TCEP for 30 minutes at room temperature then run over a Superdex 75 16/60 size exclusion column pre-equilibrated with PBS. SEC fractions containing monomeric SpyCatcher S49C were pooled, and the concentration was measured by UV absorption at 280 nm (SpyCatcher S49C $\varepsilon = 14.44\ mM^{-1}\ cm^{-1}$). Single-use aliquots were flash-frozen and stored at -80C until use.

MBP: pET28a SpyTag003-MBP was a gift from Mark Howarth (Addgene plasmid # 133450 ; http://n2t.net/addgene:133450 ; RRID:Addgene_133450). pET28-MBP-TEV was a gift from Zita Balklava & Thomas Wassmer (Addgene plasmid # 69929 ; http://n2t.net/addgene:69929 ; RRID:Addgene_69929). For expression of either SpyTag003-MBP or MBP, each plasmid was transformed into BL21 (DE3) *E. coli*. Colonies were picked and grown for 6 hours in LB + 50 ug/mL Kanamycin at 37 C. The starter culture was used to inoculate 1L flasks of autoinducing media (*2*) + 50 ug/mL kanamycin and grown overnight at 30 C. Spun down cell paste was resuspended in 350 mL of lysis buffer (50 mM $Na_2HPO_4$, 300 mM NaCl, 10 mM Imidazole) supplemented with lysozyme, benzonase, 1 mM $MgSO_4$, and protease inhibitor tablets (Roche cOmplete, EDTA-free). Cells were lysed using a tip sonicator (60% power for 2x 5 minutes, 3 seconds on, 9 seconds off) and one pass (20K PSI) through a cell disruptor. Lysates were cleared by centrifugation for 50 minutes at 40,000 RPM on a Ti45 rotor and the resulting supernatant was incubated with 1.25 mL pre-equilibrated Ni-NTA resin per liter of culture. After a two-hour incubation, Ni beads were washed in wash buffer (50 mM $Na_2HPO_4$, 300 mM NaCl, 30 mM Imidazole), and then the proteins were eluted with elution buffer (50 mM $Na_2HPO_4$, 300 mM NaCl, 350 mM Imidazole). Peak elution fractions were pooled and purified by size-exclusion chromatography with a Superdex 75 16/60 column equilibrated in PBS.

### AuNP Synthesis

Thiol-protected gold nanoparticles were synthesized as previously described (*3*). $HAuCl_4{\bullet}3H_20$ and 3-Mercaptobenzoic acid (3-MBA) were dissolved in methanol at 28 mM and 84 mM, respectively, and mixed at a 3-MBA:$AuCl_4$ molar ratio of 7:1. 2.5 volumes of water were added and the pH of the solution was adjusted to 13 with NaOH. The solution was then placed on a platform agitator at room temperature and mixed for 16 hours. The following day, the solution was diluted with methanol and water to a final concentration of 2.5 mM 3-MBA and 27% methanol. $NaBH_4$ was dissolved in cold water at 150 mM and then added to the reaction mixture at a final concentration of 2 mM. The reaction was allowed to mix on a platform agitator for 4.5 hours at room temperature, over which time the reaction mixture gradually darkens with the formation of AuNPs. The reaction product is precipitated by adding two volumes of methanol and NaCl to a final concentration of 100 mM, and centrifugation at 4,785 g (5000 RPM in a JLA/8-1 rotor) for 10 minutes. AuNPs were washed twice in 75% methanol + 100 mM NaCl, air-dried overnight in a chemical fume hood, and then resuspended in water. The concentration was by sample absorbance at 510 nm, using an AuNP extinction coefficient of $4.35x10^5$ $M^{-1}$ $cm^{-1}$ (*4*).

SpyCatcher Conjugation and Size Measurements

To generate SpyCatcher-functionalized AuNPs, 5 µM SpyCatcher S49C and 30 µM 3-MBA-protected AuNPs were mixed in PBS and allowed to react for 30 minutes at room temperature. The reaction was quenched, and the remaining AuNP conjugation sites passivated, adding reduced glutathione to a final concentration of 2.5 mM, which was allowed to react for an additional 30 minutes. After passivation, the reaction was filtered through a 0.22 µm spin filter and injected onto a Superdex 75 10/300 GL size exclusion column equilibrated in PBS. Fractions were analyzed by native PAGE using either 10% glycerol, 12% polyacrylamide gels in Tris-borate-EDTA buffer or commercial Any-KD Tris-glycine gels. Final SpyCatcher-AuNP concentrations were measured by AuNP absorbance at 510 nm, using an AuNP extinction coefficient of $4.35x10^5$ $M^{-1}$ $cm^{-1}$.

We measured the diameter of AuNPs and 1:1 SpyCatcher-AuNP conjugates using TEM and dynamic light scattering (DLS). In both cases, samples were diluted to 1 µM in PBS. For TEM, samples were frozen on copper R1.2/1.3g grids by plunge freezing in liquid ethane using a Leica EMGP plunge freezer set to 25 degrees C and 95% humidity. Single projection images were collected on a 200 keV Glacios TEM at a pixel size of 1.17 Å/px on a Gatan K2 direct electron detector. Images were collected as dose-fractionated movies at a defocus of -1.5 µm with 40 frames and a total exposure time of 1.6 seconds. For line profile analysis to measure the diameter of the AuNPs, movies were first motion-corrected with MotionCorr2 and then summed. Intensity profiles through 928 individual AuNPs were measured using FIJI/ImageJ, aligned to the peak intensity value, and averaged. DLS measurements were performed using a DynaPro NanoStar (Wyatt Technologies) and quartz cuvettes. DLS measurements were repeated three times on samples from independent SpyCatcher-AuNP conjugations that used two independent AuNP synthesis batches.

Immunocytochemistry and Confocal Microscopy

For fluorescence imaging of SpyTag-GluA2 knock-in and wild-type cultures, neurons were grown on glass coverslips as described above. SpyCatcher S49C was labeled with Janelia Fluor 646 maleimide (Tocris) following manufacturer instructions and free dye was removed Zeba dye removal spin columns. To surface stain SpyTag-GluA2 AMPARs, half the growth medium in each well was exchanged for fresh medium containing SpyCatcher-JF646 at a final concentration of 500 nM. Cultures were returned to a 37C incubator for 30 minutes for labeling, followed by two rinses with PBS + 4% sucrose and two ten-minute washes at room temperature on a plate agitator. After washing, coverslips were protected from light and fixed for 10 min in 4% PFA + 4% sucrose (in PBS) at room temperature. Following fixation, coverslips were rinsed twice in PBS, then blocked/permeabilized in PBS + 0.1% Triton X-100 + 3% BSA (TBP) for 1 h. For antigen competition experiments, coverslips were washed twice with a physiological saline solution containing (in mM) 140 NaCl, 5 KCl, 10 Glucose, 10 HEPES-NaOH, and 3% BSA (pH 7.4, ~310 mOsm) and then stained with SpyCatcher-AuNPs diluted in the same buffer at a final concentration of 250 nM. Neurons were labeled for 30 minutes in a 37-degree incubator. Following AuNP labeling, neurons were rinsed twice in a physiological saline solution and then returned to their original culture growth medium. Half the medium was exchanged with fresh growth medium containing SpyCatcher-JF646 at a final concentration of 500 nM, and the remainder of the fluorescence labeling was conducted as described above.

Fixed and permeabilized coverslips were stained overnight at 4°C in primary antibodies diluted in TBP. The following primary antibodies were used: rabbit polyclonal anti-GluA2 (1:500, Synaptic Systems Cat# 182 103, RRID:AB_2113732), mouse monoclonal anti-PSD-95 clone K28/43 (1:500, Antibodies Incorporated Cat# 75-028, RRID:AB_2292909), rabbit polyclonal anti-synaptophysin (1:500, Synaptic Systems Cat# 101 002, RRID:AB_887905), mouse monoclonal anti-MAP2 clone 198A5 (1:500, Synaptic Systems Cat# 188 011BT, RRID:AB_11042001). After staining with primary antibody, coverslips were rinsed twice in TBP and then washed 3-4 times for 5 minutes in TBP on a platform agitator at room temperature. Secondary antibody staining was done for 1 hour at room temperature using at 1:500 dilutions of antibody in TBP. The following secondary antibodies were used: goat polyclonal anti-mouse IgG2a Alexa Fluor 488 (Thermo Fisher Scientific Cat# A-21131, RRID:AB_2535771), goat polyclonal anti-mouse IgG1 Alexa Fluor 555 (Thermo Fisher Scientific Cat# A-21127, RRID:AB_2535769), goal polyclonal anti-rabbit IgG Alexa Fluor 488 (Thermo Fisher Scientific Cat# A-11008, RRID:AB_143165). After secondary antibody staining, coverslips were rinsed twice in TBP and then washed 3-4 times for 5 minutes in TBP on a platform agitator at room temperature. Coverslips were fixed again for 10 min in 4% PFA + 4% sucrose in PBS at room temperature, followed by two rinses in PBS and a final rinse in $ddH_20$. Coverslips were air dried while protected from light and then mounted on glass slides using ProLong Diamond antifade mountant. Mounted coverslips were allowed to cure, protected from light at room temperature for 12 hours, then stored at 4C until imaging.

Confocal images were acquired on a Leica SP8 inverted confocal microscope with identical acquisition settings applied to all samples within an experiment. 2048x2048 images (60 nm/px) were collected using a 1.4 NA 63X oil-immersion objective. Multi-channel images were adjusted for chromatic aberration using a rigid body transformation calculated from images of 100 nm Tetraspeck beads. Representative images were linearly adjusted for brightness and contrast to facilitate visual inspection. All such changes were applied after analysis and were made identically for all experimental conditions.

For image intensity quantification, all images were first background subtracted using a rolling-ball method with a width of 20 pixels. For line-scan analysis, a 30 µm x 10 µm rectangular section of dendrite was selected for analysis based on the MAP2 signal, and lines were drawn through individual synapse puncta identified in either the GluA2, PSD-95, or synaptophysin channel. Intensity values were normalized to the minimum and maximum values along the scan line for each channel. For quantification of SpyCatcher-JF646 intensity, the GluA2 channel was used to generate a mask using thresholding by the Otsu method in FIJI/ImageJ. The 'Analyze Particles' function in ImageJ was used to generate punctate regions-of-interest (ROI) for all GluA2 puncta in the mask image. SpyCatcher-JF646 signal was quantified within these puncta as integrated density (*i.e.*, the sum of all pixel values in a puncta) and normalized to the integrated density of GluA2 for the same ROI. To quantify the degree of the block by antigen competition with SpyCatcher-AuNPs, the signal intensity in wild-type cultures was taken as the noise floor and subtracted from the mean of the knock-in and antigen-competition knock-in conditions. The remaining signal in the antigen-competition conditions normalized to the knock-in alone condition was measured as the % block by pre-incubation with SpyCatcher-AuNPs. Analysis was conducted on images from 2 independent cultures of both SpyTag-GluA2 knock-in and wild-type animals.

Transient overexpression in HEK293T and flow cytometry staining experiments

Adherent HEK293T were seeded the day before transfection in a 6-well plate, pre-treated with poly-L-lysine, at a density of ~0.3 x 106 cells/well (25% confluency). The next day, cells were changed into fresh HEK293T media (Dulbecco's Modified Eagle Medium with 4.5 g/L D-glucose, L-glutamine, 110 mg/L sodium pyruvate, 10% FBS, 50 units/mL penicillin, and 50 µg/mL streptomycin) several hours before calcium phosphate transfection. To transiently transfect cells, a 64 µL mixture containing 6.4 µL of 2.5M calcium phosphate and 0.6 µg of pCl SpyTag-SEP-GluA2-SpyTag plasmid was added dropwise to 64 µL of 2X HEPES (280 mM NaCl, 10mM KCl, 1.5 mM $Na_2HPO_4$, 12 mM dextrose, 50 mM HEPES) while being gently vortexed. This reaction mixture was incubated for 1 minute, vortexed vigorously for 15-30 seconds, and added to cells dropwise. The next day, the media was exchanged, and the cells were used for staining experiments 48 hrs after initial transfection. Wild-type HEK293T condition cells were mock transfected as described above, but water was added instead of plasmid.

For flow cytometry staining experiments, cells were detached by aspirating out media and incubating with PBS (without Ca2+ or Mg2+) supplemented with 10 mM EDTA for 10 minutes at 37C. Cells were spun down and washed with PBS containing 2.5% BSA twice. Cells were incubated with 50 nM SpyCatcher-AlexaFluor647 for 5 minutes, washed twice, and resuspended in cold wash buffer until analyzed on a BD Accuri C6 Plus flow cytometer. Single cells were gated and analyzed for APC (AlexaFluor647) and FITC (SEP) signals using FCS Express 7 or FloJo software.

CryoEM Sample Preparation

Samples were vitrified by plunge freezing using a Leica EMGP plunge freezer set to 25 degrees C and 95% humidity. Before plunge freezing, neurons cultured on EM grids were labeled with SpyCatcher-AuNPs. Dishes containing grids were first gently washed twice with a physiological saline solution containing (in mM) 140 NaCl, 5 KCl, 10 Glucose, 10 HEPES-NaOH, and 3% BSA (pH 7.4, ~310 mOsm). SpyCatcher-AuNP was diluted into the same buffer at a final concentration of 250 nM, mixed thoroughly, and then added to each glass bottom dish, replacing the wash buffer. Neurons were then returned to a 37-degree incubator for 30 minutes to allow labeling to occur. During this labeling period, the plunge freezer was cooled to liquid nitrogen temperature, and the humidity chamber was equilibrated to minimize the delay between AuNP labeling and freezing. After the 30-minute labeling incubation, culture dishes were gently rinsed twice with buffer, then washed 2-3 times for 10 minutes at 37 C, with buffer exchanges between each wash. During the final wash, the samples were brought to the plunge-freezing area, and liquid ethane was condensed for plunge-freezing. The total time between removing the SpyCatcher-AuNP labeling solution and plunge freezing was, therefore, 30-45 minutes including wash steps and transport to the plunge-freezer. Grids were blotted from the back side for 5 seconds, then immediately plunged into liquid ethane. Vitrified grids were clipped into cryo-FIB autogrids for FIB milling and TEM tilt series data collection.

Cryo-FIB Milling

Vitrified neurons were loaded into an Aquilos 2 cryo-FIB-SEM cooled to -190 C in a 35-degree pre-tilt shuttle, and lamellae were prepared as previously described (*1*). Initial platinum sputter coating was done for 15 seconds at 30 mA and 0.10 mBarr, followed by GIS coating for 15 seconds with the GIS needle perpendicular to the grid plane. Regions of interest were identified by SEM. AutoTEM software was used to prepare lamellae at a 9-degree milling angle with the following settings: rough milling – 0.3 nA, 3 µm pattern separation in Y using rectangle patterns; medium milling – 0.1 nA, 1 µm pattern separation using cleaning-cross sections and 1-degree over/under-tilt; fine milling – 50 pA, 400 nm pattern separation using cleaning-cross sections and 0.5-degree over/under-tilt; polishing – 30 pA, 175 nm pattern separation using rectangle patterns and performed manually. Endpoint monitoring was performed using SEM scans of the lamella at 3 keV with a loss of charging contrast or breaking of the GIS layer taken as a stopping point. Final lamellae were sputter coated with platinum for 10 seconds at 7.0 mA and 0.10 mBarr immediately before unloading and storage.

Cryo-EM Data Collection

Milled grids were loaded onto Titan Krios TEM microscopes equipped with either a Gatan energy filter and Gatan K3 direct electron detector (8 tomograms) or a SelectrisX energy filter and Falcon-4i direct detector (14 tomograms). For the Gatan K3, tilt series data were collected using SerialEM in low-dose mode at a physical pixel size of 1.735 Å/px. The energy filter slit width was set to 20 eV. For the Falcon-4i, data were collected in low-dose mode using Tomo5 software and a pixel size of either 1.59 or 1.69 Å/px. The energy filter slit width was 10 eV. In all cases, tilt series were collected in three-degree increments to +/- 60 degrees, starting at a 9-degree tilt to offset the tilt of the lamellae using a dose-symmetric tilt scheme with a grouping of two (tilt sign inversions every other step). The dose per tilt was 3.2 e/Å$^2$, resulting in a total dose of 131.2 e/Å2 over 41 tilts. Images were collected as dose-fractionated movies with a per-frame dose of 0.21 e/Å$^2$.

Tomogram Reconstruction and Processing

Motion and gain correction was performed in MotionCorr2, followed by initial CTF estimation in Warp. Tilt series stacks were first aligned using AreTomo to evaluate whether they contained synapses and assess overall quality. Tilt series were discarded if they did not include synapses – marked by the presence of presynaptic vesicles, a ~25 nm width synaptic cleft, and a PSD – or if they did not reach a minimum of +/- 45 degrees. Select tomograms were subsequently aligned in IMOD using AuNPs as fiducial markers. Aligned stacks were binned by eight to a pixel size of 1.36 nm, and tomograms were reconstructed in IMOD using weighted back-projection.

For segmentations, tomograms were reconstructed in IMOD at bin 8, corresponding to pixel sizes of 13.88, 12.72, or 13.52 Å/px, using weighted back-projection and processed using Isonet (*5*) to compensate for missing wedge effects and perform denoising. Isonet-processed bin-8 tomograms were used for automated membrane segmentation using MemBrain-seg (*6*), and the resulting segmentations were manually corrected in Amira. Segmentation of pre- and postsynaptic membranes was performed as previously described (*1*). A distance field from each membrane was calculated, and membrane voxels greater than 10 nm and less than 40 nm from the opposing membrane were considered the PSD and active zone membranes, with the extracellular space between them defining the synaptic cleft. The PSD region was defined as the postsynaptic intracellular space less than 100 nm from the PSD membrane. PSD protein density was defined as voxels 1.5 standard deviations above the mean voxel intensity value of the PSD region that was in continuous contact (either directly or indirectly) with the PSD membrane. This additional contact criteria effectively removed other proteins passing through the PSD region, such as actin filaments that did not contact the membrane. Membrane proximal synaptic vesicles were segmented manually in Amira, and their center coordinates were picked in IMOD.

For localizing AuNPs, tomograms were reconstructed in IMOD at bin 2, corresponding to pixel sizes of 3.47, 3.18, or 3.38 Å/px, using weighted back-projection. AuNPs coordinates were picked using Etomo's findbeads3d program, a size of 8 pixels, a minimum spacing between localizations of 7 pixels, and a correlation threshold value of 0.75. Minimum and maximum z-slice values were also provided to exclude a region of ~10 nm from the top and bottom surfaces of each lamella. This effectively excluded prominent surface contaminants and the platinum sputter coat from being counted as AuNPs. All AuNP picks in each tomogram were manually inspected after automated picking. AuNP density was calculated as the number of localized AuNPs in a synapse divided by the PSD membrane area for that synapse.

Spatial Statistics

AuNP coordinates from each tomogram were used to analyze the spatial distribution of particles using custom scripts written in Python. As a control, for each tomogram, the same number of localizations were randomly placed into the synaptic cleft volume with a minimum distance of 2.5 nm, matching the minimum localization spacing from the observed data. 50 randomized point patterns were generated for each tomogram. Nearest-neighbor distances (NND) were measured as 3D Euclidean distances in nanometers. Histograms of pooled NND values for all tomograms were fit with a Gaussian mixture model with four components.

Univariate Ripley K functions, $\widehat{K}(r)_{Univ.}$, measure deviations from spatial homogeneity in point patterns and were calculated as:

$$\widehat{K}(r)_{Univ.} = \lambda^{-1} \sum_{i \neq j} k_{ij}^{-1} \frac{I(d_{ij} < r)}{N}$$

where r is the test distance, $d_{ij}$ is the Euclidean distance between the i-th and j-th particle, I($d_{ij}$ < r) is a counter function that returns 1 if $d_{ij}$ is less than r and 0 if not, N is the total number of particles in the test volume (the synaptic cleft), $k_{ij}$ is an edge correction factor equivalent to the fraction of the volume of a sphere of radius $d_{ij}$ contained within the test volume, and $\lambda$ is the overall particle density (number per unit volume). K functions can be mean and variance normalized such that point patterns reflecting complete spatial randomness fall along the 0 line. The mean and variance normalized form, sometimes termed H functions, $\widehat{H}(r)_{Univ.}$, are calculated in 3D as:

$$\widehat{H}(r)_{Univ.} = \sqrt[3]{\frac{3}{4}\frac{\widehat{K}(r)}{\pi}} - r$$

H functions for each individual tomogram were computed in this manner. For population analysis, samples were pooled according to the total number of particles per tomogram:

$$\widehat{H} = \frac{\sum_i N_i H_i}{\sum_i N_i}$$

Simulation envelopes were calculated by pooling all simulations for each tomogram (1100 total = 50 simulations x 22 tomograms). Statistical significance was determined by maximum absolute deviation (MAD) testing.

For bivariate analysis of the point patterns of both AuNPs and membrane-proximal synaptic vesicles, a bivariate extension of the K/H function was used.

$$\widehat{K}(r)_{Ves,AuNP} = \sum_{Ves_i} \sum_{AuNP_j} k^{-1}_{Ves_i AuNP_j} \frac{I(d_{Ves_i AuNP_j} < r)}{\lambda_{Ves} \lambda_{AuNP} V}$$

$$\widehat{H}(r)_{Biv.} = \sqrt[3]{\frac{3}{4} \frac{\widehat{K}(r)_{Ves,AuNP}}{\pi}} - r$$

Where $Ves_i$ is the i-th vesicle localization and $AuNP_j$ is the j-th AuNP localization, I is a counter function as above which returns one if the distance between a vesicle and AuNP is less than the test distance and zero otherwise, $\lambda_{ves}$ and $\lambda_{AuNP}$ are the total densities of vesicles and AuNPs per unit volume, V is the volume of the synaptic cleft, and $k_{Ves_i AuNP_j}$ is an edge correction factor.

Control data for bivariate analysis was generated by randomly reassigning the labels of synaptic vesicle and AuNP points to preserve the overall structure of the collective point pattern. This shuffling was repeated 110 times per synapse, and individual H functions were computed using MAD tests. To pool these data across synapses for population analysis, data were normalized as previously described (*7*), such that the 99% simulation envelope ranged from zero to one. Statistical significance was assessed using MAD tests, comparing the averaged H function of the observed data to all 2310 simulations (110 simulations x 21 synapses). One synapse tomogram from the dataset was excluded from bivariate analysis, since it did not have any vesicles within 10 nm of the plasma membrane.

HDBSCAN Cluster Analysis

Identification of discrete clusters of AuNP-labeled AMPARs was performed using the hierarchical density-based spatial clustering algorithm with noise (HDBSCAN) (*8*). HDBSCAN is an extension of the DBSCAN algorithm that uses hierarchical clustering to select particle clusters based on stability, allowing the selection of clusters of variable density and minimizing subjective parameter selection. Clusters contained a minimum number of 12 particles (min_cluster_size), equivalent to 6 double-labeled AMPARs. Six neighbor points within a given distance (min_samples) were required to consider a point part of a cluster. Unlike DBSCAN, which relies on a user-supplied radius value ε for distance and, therefore, sets a fixed cluster density threshold, HDBSCAN selects clusters based on cluster persistence across a range of distance values. The resulting clusters can, thus, have variable densities. Finally, clusters within

25 nm of one another were merged (cluster_selection_epsilon). Cluster area was measured as half the surface area of the convex hull of each set of clustered AuNPs. The percent of the PSD membrane area covered by clusters was calculated as the sum of all cluster areas divided by the PSD membrane area.

Paired Cross-Correlation Function Analysis

To compare the spatial relationship between either AuNP-labeled AMPAR localizations or vesicle positions versus the local protein density of the PSD scaffolding complex, we computed paired cross-correlation functions similar to the previously described analysis of fluorescence single-molecule data (*9*, *10*). This analysis was performed in 2D in the plane of the synaptic cleft since our analysis was intended to focus on the lateral correlation between the two density maps (*i.e.,* spatial correlations in the membrane plane). Computing the paired-cross correlation in 3D resulted in a prominent peak in the paired cross-correlation across all data – observed and simulated – that reflected increased correlation in the pre-to-postsynaptic axis due to PSD protein density being concentrated at the membrane vs further into the cytoplasm. Boundary-corrected local density maps of the PSD were first generated in 3D in Amira using a window of 11x11x11 voxels (~14 nm per side). To generate density maps from AuNP and vesicle coordinates, AuNP and vesicle coordinates were converted to binary 3D volumes matching the pixel size and dimensions of the PSD density maps. The best-fit plane to either the AuNP or the vesicle point pattern was used to determine the orientation of the synaptic cleft. Voxel data was projected onto this plane, and the basis was changed to orient the projections in the xy plane. AuNP and vesicle coordinate maps were filtered with a Gaussian kernel with a standard deviation of 3 voxels (for AuNPs) or 10 voxels (for vesicles). These values were chosen to reflect the approximate width of single AMPARs and vesicles, respectively. Paired cross-correlation functions, $g_c$(r), normalized to account for correlations due to the shape of the measurement region, were computed as:

$$g_c(\vec{r}) = Re\left\{\frac{FFT^{-1}(FFT(I_{AuNP}) \times conj[FFT(I_{PSD})])}{\rho_{AuNP}\ \rho_{PSD}\ FFT^{-1}(FFT(W_{AuNP}) \times conj[FFT(W_{PSD})])}\right\}$$

Where $I_{AuNP}$ and $I_{PSD}$ are the density maps of the AuNPs and PSD, $W_{AuNP}$ and $W_{PSD}$ are window regions representing the synaptic cleft volume and PSD region, respectively, with values of one inside the window and zero outside, $\rho_{AuNP}$ and $\rho_{PSD}$ are the overall density of each map within the window region, conj[] refers to the complex conjugate, and Re{} refers to the real component. This output was averaged over radial bins to give $g_c(r)$. For vesicles, the window region was calculated as the convex hull of all vesicle localizations.

Observed AuNP localization data was compared to control data generated by randomly placing the same number of particles into each synaptic cleft volume with a minimum inter-particle distance of 2.5 nm for AuNPs and 45 nm for vesicles. Fifty randomized simulations were performed per tomogram. To compute simulation envelopes, one simulation was randomly picked for each tomogram, and the average was calculated across all tomograms. This process was repeated 100 times, with 1/50 simulations selected randomly from each tomogram for each permutation.

Modeling of SpyCatcher/Tag-GluA2 AMPARs

Modeling of SpyTag-GluA2 AMPARs and covalently bound SpyCatcher-AuNPs was performed using 20 previously published cryoEM models of full-length GluA2-containing AMPARs in different states and with reported FSC resolutions less than 7Å (PDBs: 5WEM, 5WEN, 5WEO, 6NJL, 6NJM, 6NJN, 6QKZ, 7RZ4, 7RZ6, 7RZ7, 7RZ9, 7RZA, 7OCA, 7LDD, 7LDE, 8SSA, 8SS6, 8SS8, 8VJ7, 8VJ6; see Table S1). The 'Build Structure' tool in ChimeraX was used to join the GluA2 subunits in positions B and D of each model to a model of the SpyTag/Catcher complex (PDB 4MLI). The C-terminal threonine residue of SpyTag (mutated to Tyr in SpyTag003) was covalently linked to Asn25 of GluA2 with a C-N length of 1.33Å, an ω angle of 180°, and ϕ angle of -120°. Serine 49 in SpyCatcher was mutated to Cys using the 'swapaa' command with Chi angle of 67.8°. AuNPs were modeled as ChimeraX markers with a radius of 1.25 nm placed at C49 of each modeled SpyCatcher. The 'distance' command in Chimerax was used to measure the distance between residue 49 of the B/D subunits and between the modeled AuNP markers. Depending on the orientation of SpyCatcher, these two values differed by up to 1.64 nm, though on average, there was only a difference of 0.135 nm across all 20 models.

Subtomogram Averaging (STA)

The 8-tomogram dataset collected at 1.735 Å/px on a Gatan K3 direct electron detector was used for STA. Movies were reprocessed using Warp2.0 for Linux (*11*) to perform motion correction, initial CTF estimation, and tomogram reconstruction using previously generated tilt series alignments. AuNP coordinates were used as reference points to manually pick visible receptor density from IsoNet (*5*) corrected tomograms. Manual picking was limited to receptor-like density that appeared connected to an AuNP within 7 nm. 1545 AuNP coordinates were inspected, resulting in 834 manual picks (53%) centered on the post-synaptic plasma membrane.

Two additional copies of the motion-corrected average projection images were generated. In the first, AuNP coordinates picked from each tomogram were back-projected and masked with Gaussian noise using the 'rawtiltcoords' and 'ccderaser' programs in IMOD (*12*), respectively. In the second, 3D segmentations of the plasma membrane, generated using MemBrain-seg 10/21/24 6:45:00 PM, were back-projected to match the original tilt series of projection images and to create a tilt series consisting of weighted membrane masks with higher values corresponding to a greater number of segmented membrane voxels along the projection line. These weighted mask values were then normalized across the tilt series mask by dividing by the maximum value for the whole tilt series. The mean pixel intensity value was calculated for each projection image in the original tilt series data, and pixels covered by the weighted membrane mask were scaled by subtracting the difference between each observed intensity pixel and the tilt-image mean, scaled by the value of the weighted membrane mask with the following formula:

$$Pixel_{supp.} = Pixel_m - \left(Pixel_m - Pixel_{avg}\right) \times \left(Mask_m \div Mask_{max}\right)$$

Where $Pixel_m$ is the intensity value of a masked pixel, $Pixel_{avg}$ is the average intensity value of a pixel in a specific tilt-image, $Mask_m$ is the corresponding weighted membrane mask value for the pixel, and $Mask_{max}$ is the maximum mask value across the entire tilt series. The result is that pixels containing membrane had their values pushed toward the mean of the respective tilt-image, with projections along the plane of the membrane resulting in greater signal suppression compared to projections perpendicular to the membrane plane. This membrane signal suppression was performed in addition to AuNP signal subtraction.

For STA of the extracellular AMPAR domains, sub-tomograms were extracted as 3D volumes at a pixel size of 8 Å/px and a box size of 46 using WarpTools, initially from the AuNP-subtracted tilt series. An initial round of 3D refinement in Relion4 (*13*) was performed using a flat cylinder (radius 18 nm, height 7 nm) as a reference, allowing rotations but restricting translations to align all particles with the membrane in the xy plane. This resulted in an STA map showing an extracellular receptor density and a high signal-intensity PSD region. ArtiaX (*14*) was used to map back the STA map into the tomograms and to manually inspect each tomogram's particles. Particles with flipped psi angles (receptors pointed to the PSD side) or with tilt and psi angles beyond two standard deviations from the mean tilt/psi angle of all particles in the tomogram were re-oriented to match the mean. Another round of refinement was then performed, limiting tilt and psi angles to 2 s.d. and increasing the translational searches to 4 pixels. This was followed by removing duplicate particles with a threshold of 8 nm inter-particle distance, resulting in a dataset of 613 particles. Such duplicates occurred by accidental picking errors or shifts during initial 3D refinements. These particles were then reextracted from the membrane suppressed, AuNP subtracted tilt series. Three rounds of refinement were performed, limiting tilt and psi angles to 2 s.d. The first used a standard spherical mask and no reference. The second used a soft cylinder mask covering the extracellular region and transmembrane domain of the receptor density and the results from the previous refinement as a reference. The final round used a soft cylinder mask covering just the extracellular region and the previous refinement results as a reference. Finally, sub-tomograms were re-extracted from the original tilt series, including AuNPs and membranes. Refinements were performed either with or without applying C2 symmetry using the previous maps as a reference and limiting translations to 1 pixel and rotations to 1 s.d., resulting in maps with global resolutions of 31 Å as assessed by gold-standard FSC at a cutoff of 0.143.

For STA of the PSD, receptor positions and angles from AMPAR STA were used to re-extract sub-tomograms centered inside the PSD directly underneath the receptor, such that the LBD, but not the NTD of the AMPAR was contained within the extracted 3D volumes (8 Å/px, box size 46). By transferring the positions and angles from the AMPAR extracellular domain STA, we ensured that the resulting PSD STA maps were closely correlated with the AMPAR STA maps. An initial round of 3D refinement was performed without a reference and using a standard spherical mask, limiting translations to 3 pixels and tilt and psi rotations to 2 s.d., resulting in a map with several prominent intracellular protein densities and an extracellular density corresponding to the AMPAR LBD region. Following this initial refinement, 3D classification without alignment was performed using ten classes. Two prominent densities were apparent: a globular density and an elongated density, both in apparent alignment with the AMPAR extracellular region. Classes with either of these features were selected, resulting in a dataset of 460 particles (74% of the initial AMPAR particles). These particles were used for a subsequent round of refinement, using the previous map as a reference. Translations were limited to 1 pixel and rotations on all angles were limited to 2 s.d. This resulted in a final map with a global resolution of 28 Å as assessed by gold-standard FSC at a cutoff of 0.143.

Composite Model Building and Back-Plotting

The maps of the C2-symmetric AMPAR and PSD were used to build a composite map of the AMPAR-PSD scaffold complex in ChimeraX (*15*). The plasma membrane of both maps was first used for alignment to correct for the difference in extraction coordinates and small shifts between the maps. Next, the structure of the natively purified GluA1/A2/A1/A2 AMPA receptors (PDB 7LDD) was used to slightly adjust the orientations of both maps more precisely. The transmembrane region of the receptor, which includes the auxiliary subunits TARP gamma-8 and CNIH2, was used to rigid body dock the model into the PSD map, where the TM regions appeared resolved between the membrane bilayers. The C2-symmetric AMPAR map was then rotated to fit to the extracellular region of the AMPAR model. SpyCatcher/Tag (PDB 4MLI) was joined to the N-terminal of the GluA2 subunits, then fit to the map density. A 2.5 nm marker representing the AuNP was placed at residue 49 of SpyCatcher. The crystal structure of the tandem PDZ1/2 domains of PSD-95 (PDB 3ZRT) was rigid body docked into the elongated PSD density. The guanylate-kinase/SH3 domain of PSD-95 (PDB 1JXO) was rigid body docked into the second globular PSD density.

ArtiaX (*14*) was used to back-plot STA maps to the original tomogram coordinate system using .star files and maps from the final rounds of refinement. Membrane-proximal synaptic vesicles, represented as 48-nm spheres, were visualized in the same map.

Statistics

Statistical significance was set at *: $p<0.05$; **: $p<0.01$; ***: $p<0.001$. The complete dataset consisted of 22 tomograms from SpyTag-GluA2 knock-in neurons collected from four independent cultures and imaging sessions. In addition, 12 wild-type synapses from two independent cultures and imaging sessions were used to assess the specificity of SpyCatcher-AuNP labeling. Comparisons of AuNP labeling density between conditions used Mann-Whitney U tests. Statistical comparisons of AuNP localization data to simulated data by paired cross-correlation or Ripley H functions were performed using maximum absolute displacement (MAD) tests. This calculates the maximum absolute difference between each trial (real data or simulated) versus the mean of all trials (data plus simulated). The p-value is calculated as the number of simulation trials with MAD values greater than that of the true data, divided by the total number of trials. All other significance testing was performed using student's t-tests.

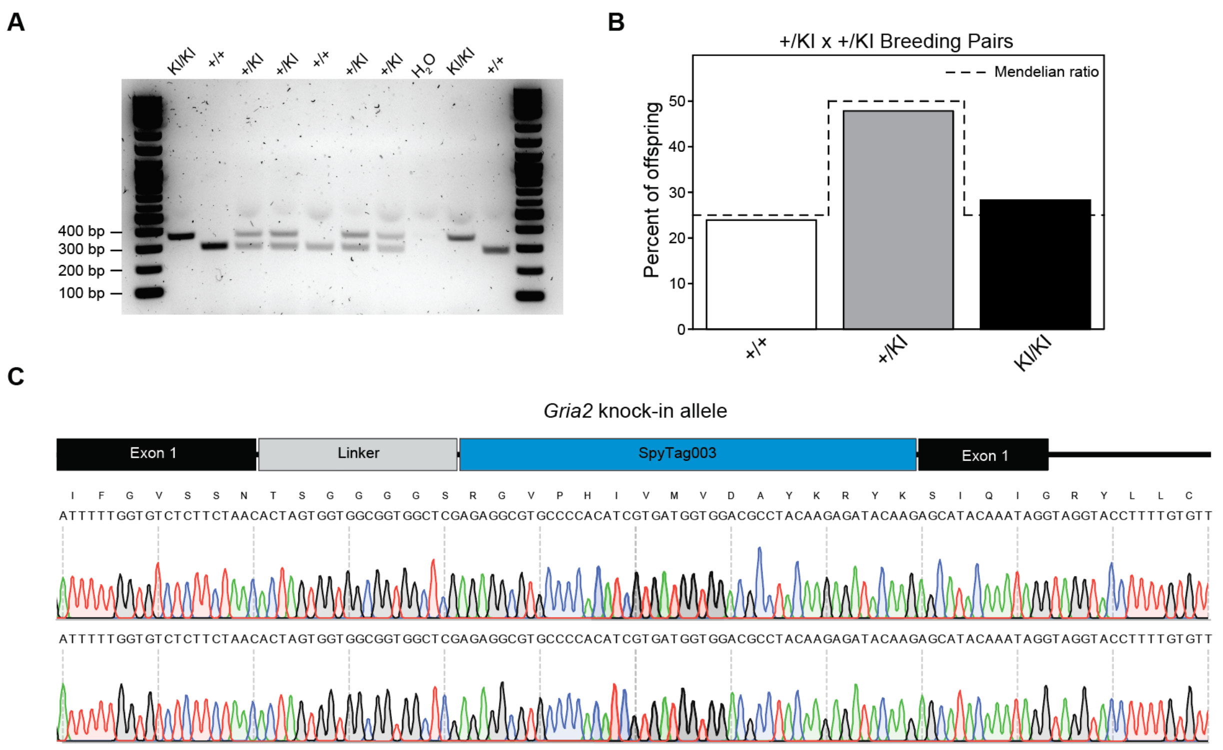


**Fig. S1. Genotyping data of SpyTag-GluA2 knock-in mice.**

**A)** Genotyping PCR gel of a 7-pup litter from a heterozygous (+/KI) breeding pair showing knock-in (371 bp) and wild-type (302 bp) bands. **B)** Percentage of offspring from +/KI breeding pairs of each genotype. Dashed lines indicate the expected Mendelian ratio (N=46 mice / 7 litters / 3 breeding pairs). **C)** Sanger sequencing results of gel purified KI/KI bands showing in-frame integration of SpyTag into the open reading from of *Gria2* exon 1. PCR primers (indicated in Fig. 1C) were outside of the donor DNA sequence.

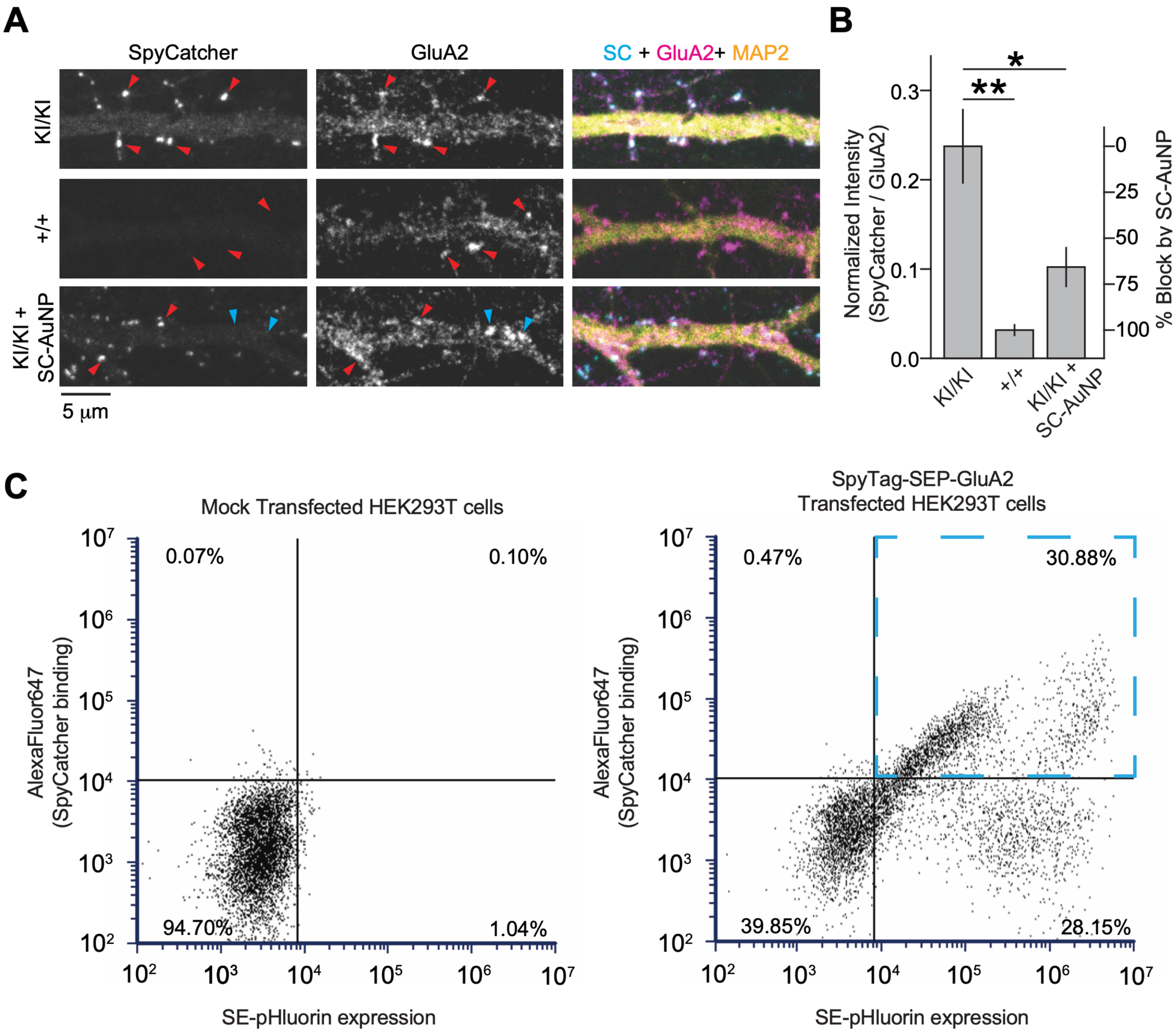


**Fig. S2. Fluorescence microscopy and flow cytometry of SpyCatcher/Tag specificity and antigen masking.**

**A)** Example images of surface staining with fluorescent SpyCatcher (left column), immunostaining with GluA2 antibodies (middle column), and merged with the dendritic marker MAP2 (right column). The top row shows staining of SpyTag-GluA2 knock-in neurons (KI/KI), the middle row control staining of wild-type neurons (+/+), and the bottom row fluorescent staining of KI/KI neurons after antigen masking with 250 nM SpyCatcher-AuNPs (SC-AuNP). Red arrowheads mark example GluA2 puncta. Blue arrowheads indicate prominent GluA2 puncta lacking fluorescent SpyCatcher staining after antigen competition. **B)** Quantification of SpyCatcher fluorescent intensity within GluA2 puncta, normalized to GluA2 intensity (N= 12 images/2 cultures KI/KI, 4 images, 2 cultures +/+, 6 images/2 cultures KI/KI + SpyCatcher-AuNP masking). The right ordinate axis indicates the signal loss by SpyCatcher-AuNP antigen masking, setting the fluorescent noise floor at the level of +/+ staining and maximum at the KI/KI staining level (**: $p<0.01$; *: $p<0.05$). **C)** Flow cytometry plots of AlexaFluor647 labeled

SpyCatcher binding to HEK293T cells, mock transfected with no plasmid (top) or HEK293T cells overexpressing SpyTag-super ecliptic pHluorin (SEP) linked GluA2 (bottom). The y-axes show AlexaFluor647 fluorescence, representing the degree of SpyCatcher-AlexaFluor647 binding to cells, and the x-axes show SEP fluorescence, representing the degree of surface-expressed SpyTag-GluA2. The Blue dashed quadrant highlights the population of SpyTag-SEP-GluA2 expressing cells binding specifically to SpyCatcher-AlexaFluor647.

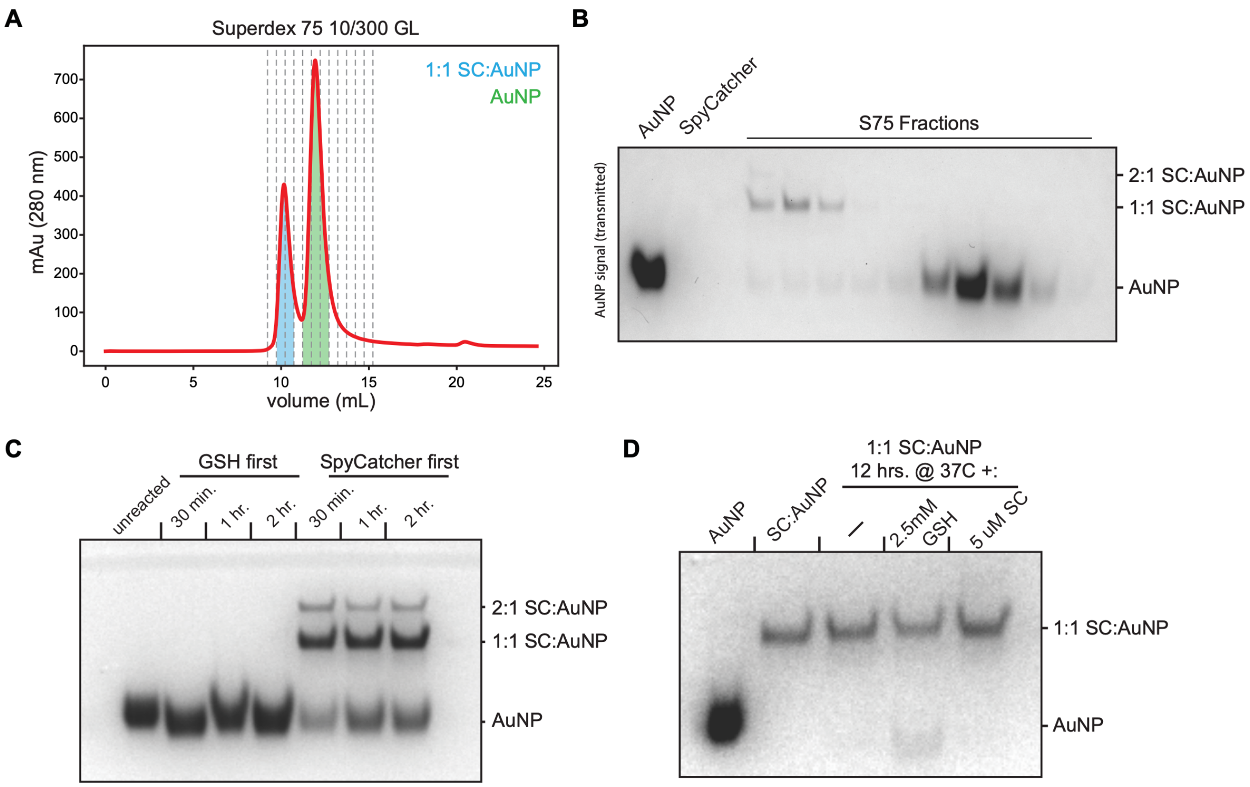


**Fig. S3. Purification and stability of SpyCatcher-functionalized AuNPs.**

**A)** Size-exclusion chromatogram of a SpyCatcher-AuNP conjugation reaction. Peaks of majority 1:1 SpyCatcher-AuNPs are highlighted in blue, and excess, unconjugated AuNPs are highlighted in green. **B)** Native PAGE of SEC fractions from a SpyCatcher-AuNP conjugation reaction. **C)** Native PAGE of representative AuNP passivation reactions with reduced glutathione (GSH). AuNPs were reacted with either GSH followed by SpyCatcher (GSH first) or SpyCatcher followed by GSH (SpyCatcher first). Reaction times are indicated above each lane. Reactions were not SEC purified. **D)** Native PAGE of GSH passivated, SEC purified 1:1 SpyCatcher-AuNPs (see Methods) incubated for an additional 12 hours at 37 C with the indicated concentrations of GSH or unconjugated SpyCatcher to assess the potential for place-exchange reactions to displace SpyCatcher or the GSH passivation.

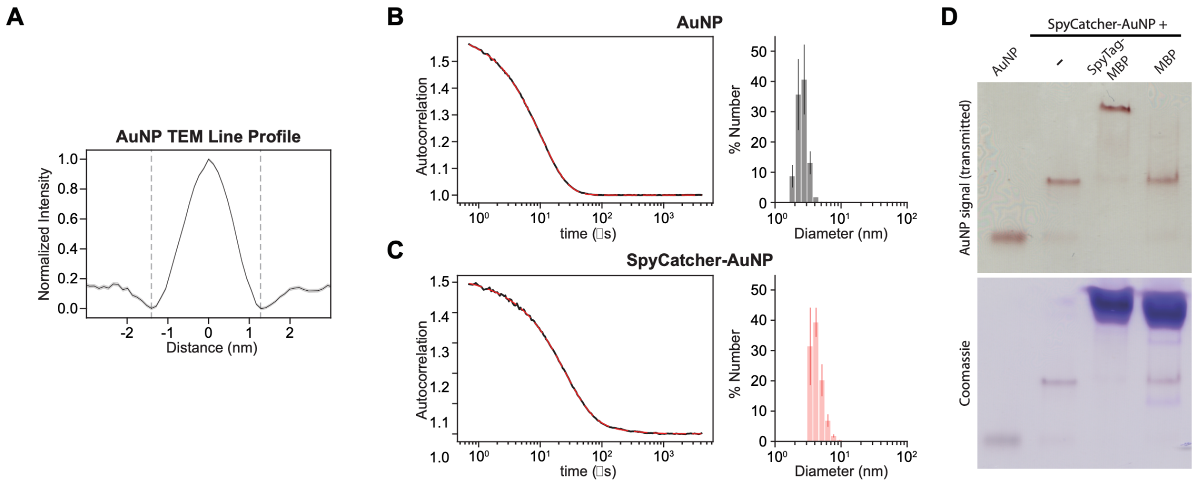


**Fig. S4. AuNP size measurement and SpyCatcher-AuNP function**

**A)** Averaged line scans through unconjugated AuNPs in cryoEM micrographs taken with 1.5 μm defocus, as shown in Fig. 2B (N=978 AuNPS). **B, C)** Representation DLS autocorrelation trace (left) and diameter histogram (right) of unconjugated AuNPs (B) and purified 1:1 SpyCatcher-AuNPs (C). The black line in the left panel is the raw autocorrelation, and the dashed red line is the model fit. Diameter histograms show the mean and standard error of three independent measurements. **D)** Native PAGE of the gel shift assay shown in Fig. 2C with the Coomassie stain of the same gel shown below.

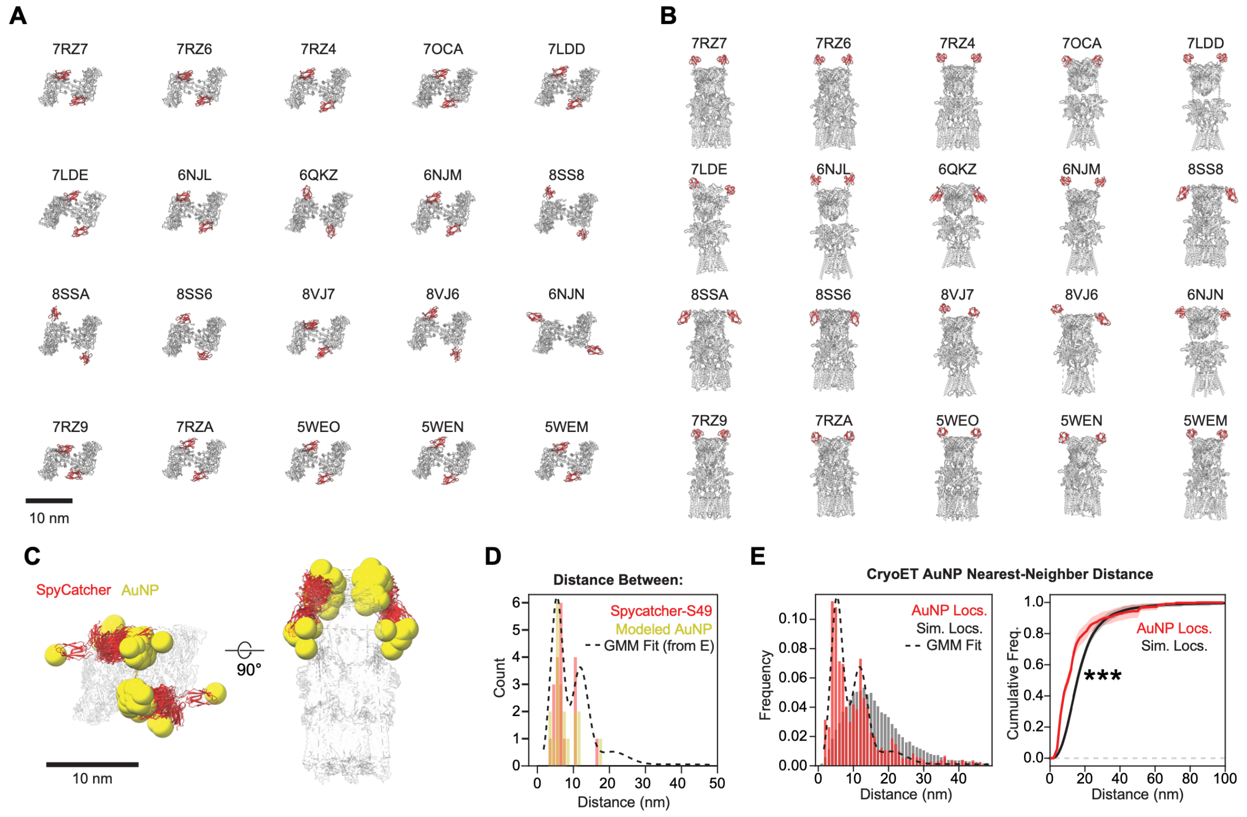


**Fig. S5. Modeling of SpyCatcher bound SpyTag-GluA2 AMPAR complexes.**

**A)** Top-down views of the N-terminal domain of GluA2-containing AMPARs (grey) joined with SpyTag/SpyCatcher (red, PDB 4MLI) at the B and D subunit positions. PDBs for each GluA2 model are indicated (see also Table S1). **B)** Side views of modeled GluA2-containing AMPARs (grey) joined with SpyTag/SpyCatcher (red). **C)** Top-down (left) and side view of superimposed modeled SpyTag-GluA2 complexes (grey, transparent) with SpyCatcher-AuNPs (red=SpyCatcher, yellow=AuNP). AuNPs are modeled as 2.5 nm spheres placed at SpyCatcher S49 (mutated to C49). **D)** Histogram of modeled distances between either SpyCatcher S49 (red) or AuNP markers (yellow). The multi-Gaussian function fit of AuNP nearest-neighbor distance values in synapse tomograms is overlayed (dashed line). **E)** Histogram (left, same as Fig. 2B) and cumulative histogram (right) showing the nearest-neighbor distance values between AuNPs in SpyTag-GluA2 knock-in synapse tomograms. The right panel shows the mean and standard error of individual synapses (N=22, red line) compared to the mean of simulated data for each synapse (N=2,200 (100 random simulations per synapse), black line). Statistical comparison was performed by MAD test (***, $p<0.001$).

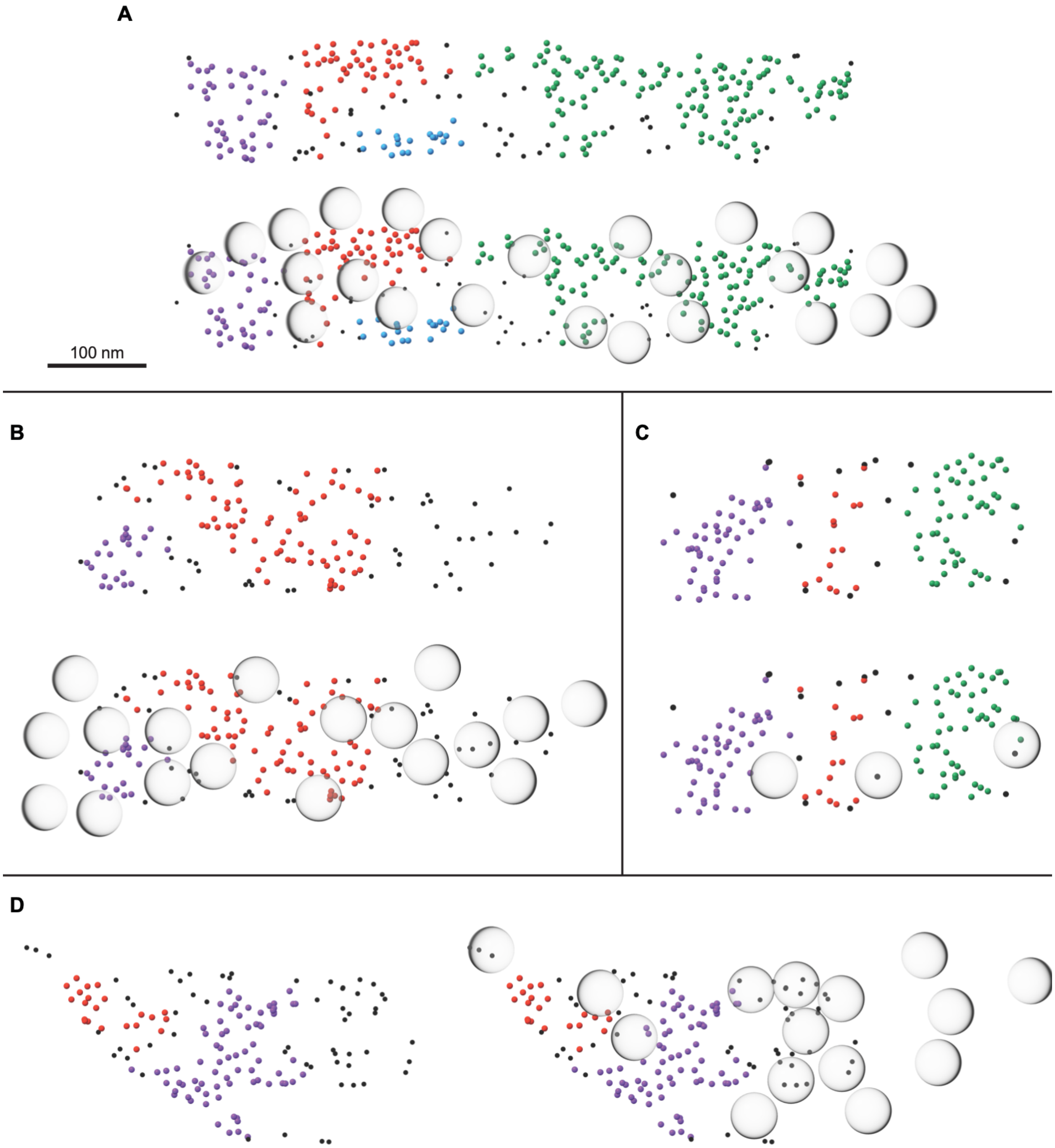


**Fig. S6. Example AuNP point patterns and overlayed membrane-proximal vesicles.**

**(A-D)** AuNP point patterns, displayed as 4 nm diameter dots, either without or with transparent overlays of all membrane-proximal vesicles. AuNPs are color-coded by HDBSCAN cluster label (green, red, purple) or black for AuNPs outside the clusters.

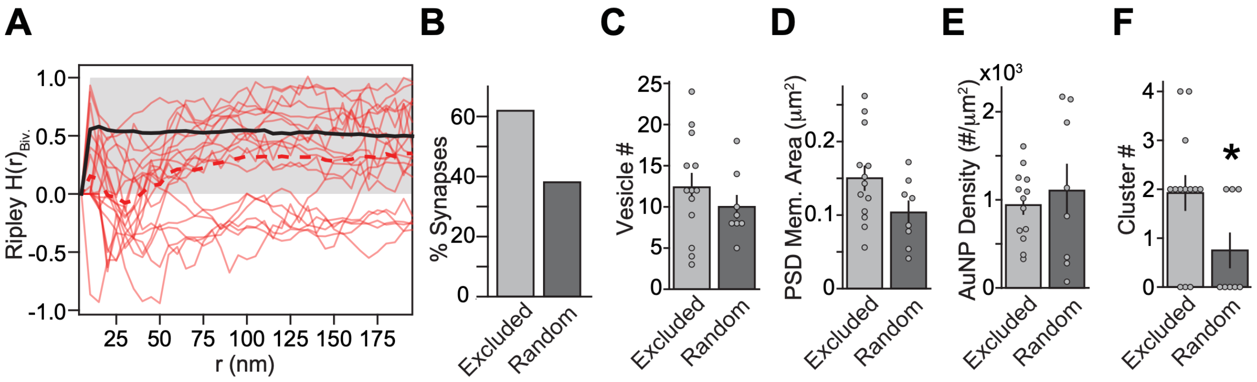


**Fig. S7. Bivariate cluster analysis of individual synapses.**

**A)** Bivariate Ripley H analysis of the relationship between synaptic vesicle position and AuNP localizations. Red traces show the value for each individual synapse, with the red dashed line indicating the mean (N=21 synapses). The black line and gray boundary show the mean of simulated data and 99% simulation envelope (N=2310), respectively. **B)** Rejection of the null hypothesis for each individual synapse. Synapses were compared to their respective randomized controls (N=110/synapse) by MAD test. Synapses that reached statistical significance ($p<0.05$) were termed "excluded" and those that did not were termed "random." The two synapse populations had no significant difference in GluA2-AMPAR density, synapse area, or membrane-proximal vesicle number. **C)** Number of membrane-proximal synaptic vesicles for excluded synapses compared to random topography synapses. **D)** PSD membrane area for excluded synapses compared to random topography synapses. **E)** Synaptic AuNP labeling density for excluded synapses compared to random topography synapses. **F)** Number of AuNP-labeled AMPAR clusters identified by HDBSCAN for excluded compared to random topography synapses. Significance was determined by student's t-test (*; $p<0.05$, N=13 excluded, 8 random).

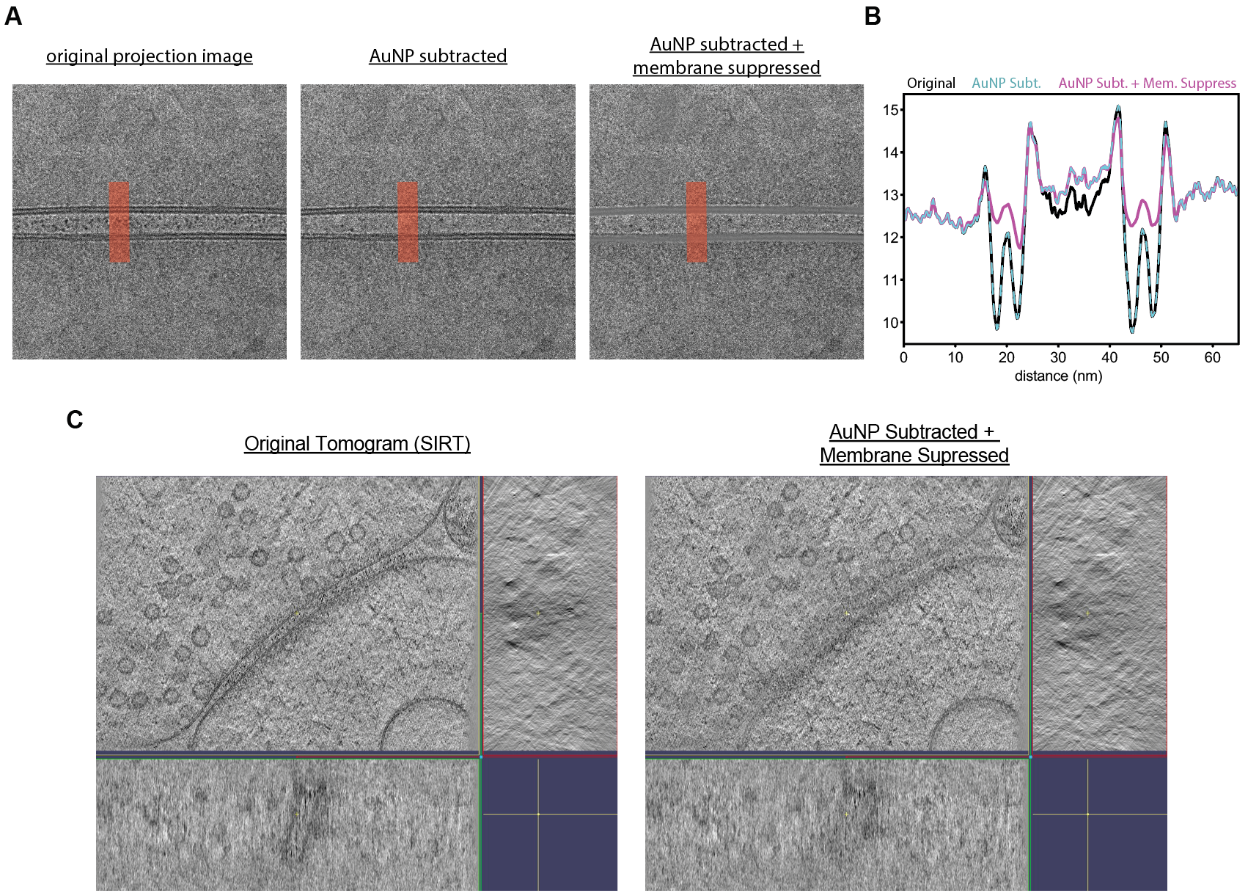


**Fig. S8. AuNP Signal Subtraction and Membrane Suppression**

**A)** Example tilt series projection images from the original stack (left), after subtraction of AuNP signal (middle), and after AuNP subtraction and membrane signal suppression (right) **B)** Line-scan profiles of the sections highlighted in red in A, showing pixel intensity values of original (black), AuNP subtracted (cyan dashed), and AuNP subtracted + membrane suppressed (magenta) projection images. **C)** Example tomogram slices (XYZ viewer) of tomograms reconstructed from the original stack (left) and after AuNP subtraction and membrane signal suppression (right).

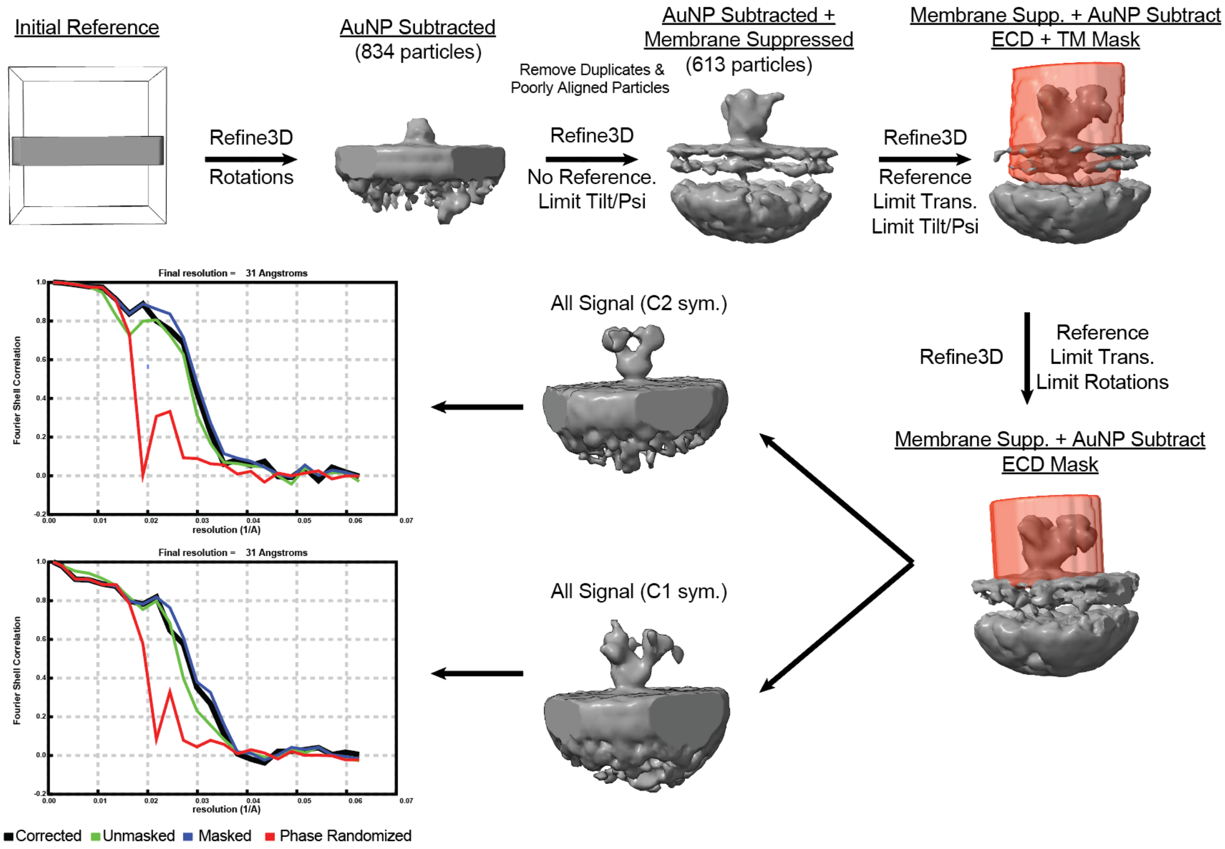


**Fig. S9. STA of AMPAR Extracellular Domains**

**A) C)** STA workflow for AMPA receptors resulting in final maps with global FSC resolution of 31 Å. Details of each refinement step are described in the methods section.

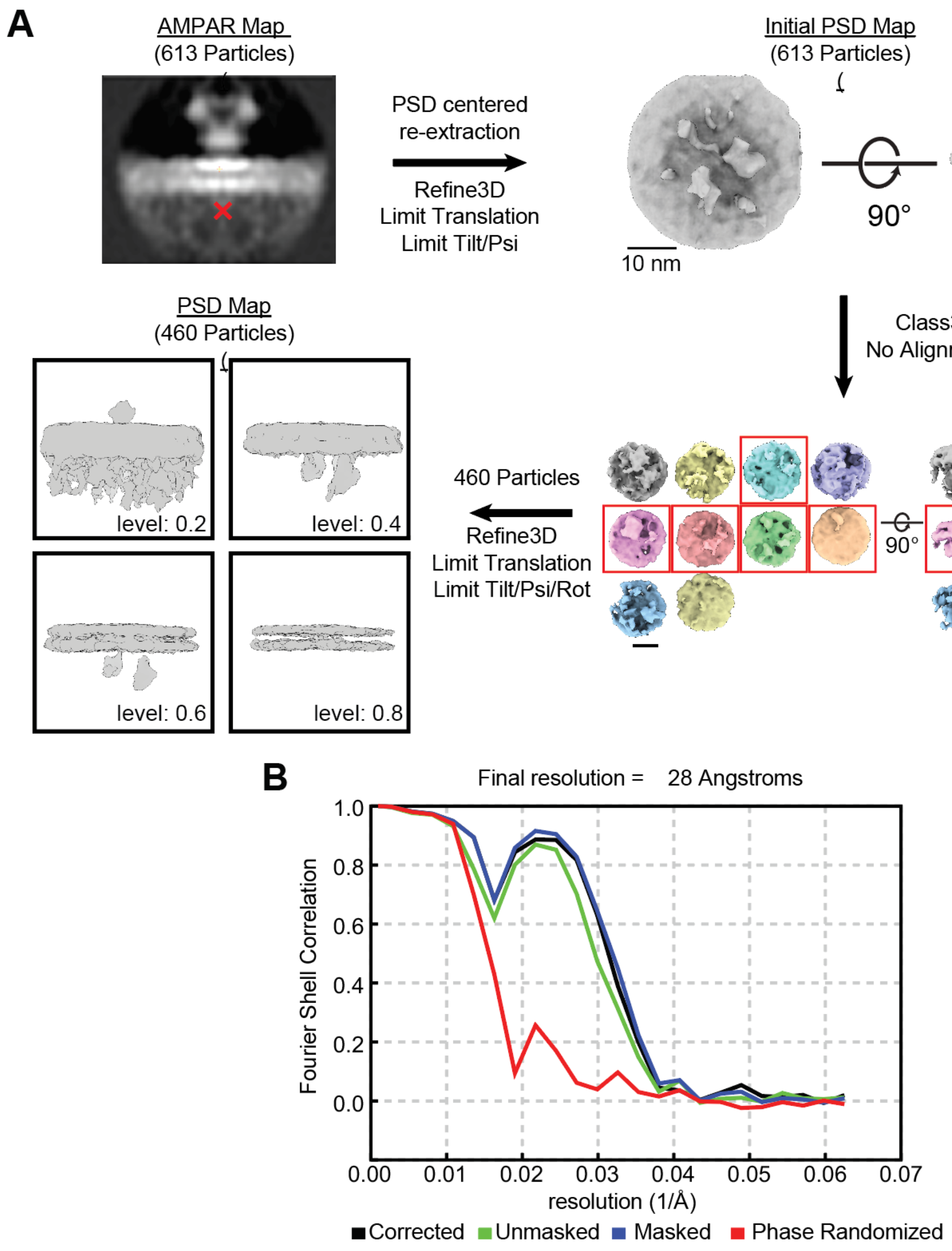


**Fig. S10. STA of AMPAR/PSD Scaffolding Complexes.**

**A)** STA workflow for AMPAR/PSD scaffolding complexes. Details of each refinement and classification step are described in the methods section. **B)** FSC curves of the final map with a reported global resolution of 28 Å.

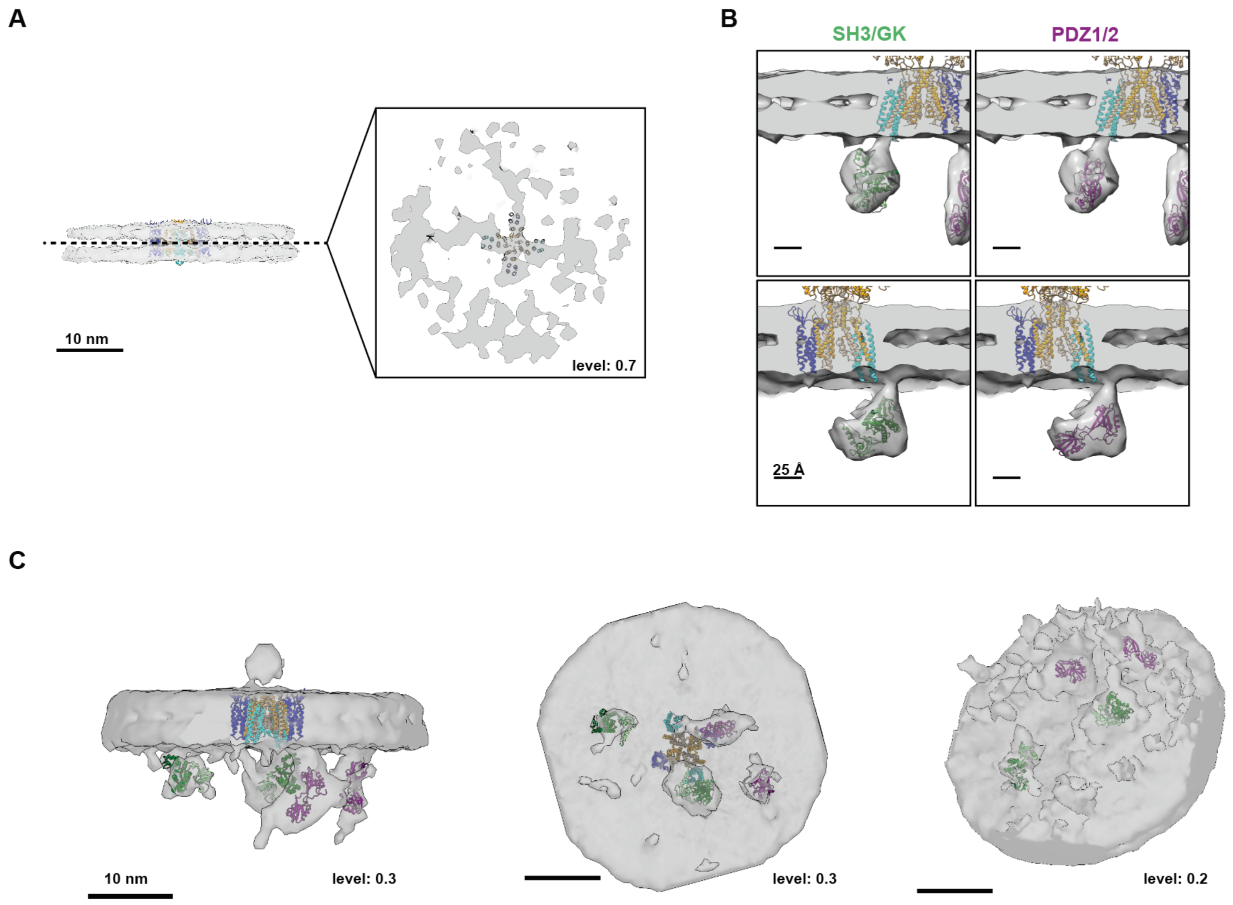


**Fig. S11. Docking of Transmembrane Helices and Alternative PSD-95 Domains**

**A)** A planar slice between the membrane leaflets of the PSD-focused STA map (surface contour level 0.7). Transmembrane portions of the docked AMPAR model (PDB 7LDD) fit a plus-shaped transmembrane density in the STA map. **B)** Comparison, showing two viewing angles of docking the crystal structures of the SH3/GK domain (PDB ID 1JXO) versus the tandem PDZ1/2 domains of PSD-95 (PDB ID 3ZRT) into the globular PSD density. **C)** Lower contour (levels indicated) views of the PSD-focused STA map showing multiple densities that can accommodate repeated copies of the folded domains of PSD-95 (PDB 1JXO, 3ZRT). Left: side view parallel to the membrane plane, middle: bottom-up view from the postsynapse, right angle view from the postsynapse.

| PDB | Pos. A/B/C/D | Aux. Subunits | Ligand(s) | Resolution (A) | Dist. SpyCatcher S49 (nm) | Dist. AuNP (nm) | State | PMID |
|---|---|---|---|---|---|---|---|---|
| 7RZ9 | A2/A2/A2/A2 | GSG1L | | 4.15 | 3.7389 | 4.2017 | closed | 34678168 |
| 7RZ7 | A2/A2/A2/A2 | TARP γ-5 | quisquilate | 4.2 | 4.33 | 3.605 | desensitized | 34678168 |
| 7RZ6 | A2/A2/A2/A2 | TARP γ-5 | glutamate | 4.4 | 4.55 | 3.46 | desensitized | 34678168 |
| 7OCA | A1/A2/A1/A2 | TARP γ-8; CNIH-2 | | 3.4 | 4.99 | 5.236 | closed | 34079129 |
| 5WEM | A2/A2/A2/A2 | GSG1L | | 6.1 | 5.125 | 5.267 | closed | 28737760 |
| 8VJ7 | A2/A2/A2/A2 | | glutamate; GYKI-52466 | 4.85 | 5.48 | 5.279 | inhibited | 38834914 |
| 7RZA | A2/A2/A2/A2 | GSG1L | quisquilate | 4.26 | 5.892 | 6.129 | desensitized | 34678168 |
| 7LDD | A1/A2/A1/A2 | TARP γ-8; CNIH-2 | ZK1 | 3.4 | 5.9207 | 5.609 | closed | 33981040 |
| 5WEO | A2/A2/A2/A2 | TARP γ-2 | glutamate, CTZ | 4.2 | 6.0407 | 6.1633 | open | 28737760 |
| 6NJL | A1/A2/A1/A2 | TARP γ-2 | ZK1 | 6.7 | 6.1033 | 5.267 | closed | 30975770 |
| 7RZ4 | A2/A2/A2/A2 | TARP γ-5 | ZK1 | 3.6 | 6.26 | 5.839 | closed | 34678168 |
| 6NJM | A3/A2/A3/A2 | TARP γ-2 | ZK1 | 6.5 | 6.432 | 6.284 | closed | 30975770 |
| 7LDE | A1/A2/A1/A2 | TARP γ-8; CNIH-2 | ZK1 | 3.9 | 6.606 | 7.06 | closed | 33981040 |
| 5WEN | A2/A2/A2/A2 | GSG1L | | 6.8 | 6.924 | 7.716 | closed | 28737760 |
| 8SS6 | A2/A2/A2/A2 | TARP γ-5; CNIH-2 | ZK1; perampanel; spermidine | 3.01 | 7.082 | 8.012 | closed | 37653241 |
| 8VJ6 | A2/A2/A2/A2 | | glutamate; GYKI-52466 | 3.5 | 10.14 | 10.654 | inhibited | 38834914 |
| 8SS8 | A2/A2/A2/A2 | TARP γ-5 | ZK1; perampanel | 2.81 | 10.56 | 11.254 | closed | 37653241 |
| 6QKZ | A1/A2/A1/A2 | TARP γ-8 | NBQX | 6.3 | 10.584 | 10.09 | closed | 30872532 |
| 8SSA | A2/A2/A2/A2 | TARP γ-5; CNIH-2 | glutamate; spermidine | 3.88 | 10.82 | 11.525 | desensitized | 37653241 |
| 6NJN | A1/A2/A3/A2 | TARP γ-2 | ZK1 | 6.5 | 16.13 | 17.77 | closed | 30975770 |

**Table S1. Model information for SpyTag-GluA2 AMPAR models**

Published structures of GluA2-containing AMPARs used to model predicted AuNP nearest-neighbor distances. The table is sorted by the measured distance between SpyCatcher residue S49 labeling GluA2 in positions B/D. Red highlighted rows indicate structures where predicted distances fall into the second peak of the measured nearest-neighbor distance histogram (see Supp. Fig. S4D)

**Movie S1. Composite Map and Model of the AMPAR – PSD-95 Scaffolding Complex**

Spin movie showing the composite STA maps of the AMPAR extracellular domains (AMPA-ECD map, teal) and the PSD scaffolding complexes (PSD map, grey). At the halfway point, the docked models of the native AMPAR (PDB-7LDD: GluA1/A2, TARP-γ8, CNIH2), SpyCatcher/Tag (PDB-4MLI), and the PDZ1/2 tandem (PDB-3ZRT) and SH3/GK domains (PDB-1JXO) of PSD-95 are shown, with color coded labels.

**Movie S2. Visualization of AMPAR – PSD Scaffolding Complex Topography in a Tomogram.**

A movie showing sequential XY slices through a tomogram, denoised with cryoCARE (*16*). The STA maps of the AMPAR-ECD (orange) and PSD scaffolds (purple) are plotted in their position and orientation as determined by STA. Synaptic vesicle positions (cyan) and the postsynaptic plasma membrane (grey) are also shown as 3D volumes. At the 45 second mark, the color code changes to display AMPARs that are anchored to PSD scaffolds in orange (defined as those with PSD STA maps that were not removed during 3D classification) and free AMPARs in green.
